# Rare-variant aggregate association analysis using imputed data is a powerful approach

**DOI:** 10.1101/2025.04.17.649394

**Authors:** Tianyi Liu, Paul L. Auer, Gao T. Wang, Elnaz Naderi, Andrew T. DeWan, Suzanne M. Leal

**Affiliations:** Center for Statistical Genetic, Gertrude H. Sergievsky Center, Department of Neurology, Columbia University Medical Center, New York, NY, USA; Division of Biostatistics, Data Science Institute, and Cancer Center, Medical College of Wisconsin, Milwaukee, WI, USA; Department of Chronic Disease Epidemiology and Center for Perinatal, Pediatric and Environmental Epidemiology, Yale School of Public Health, New Haven, CT, USA; Taub Institute for Alzheimer’s Disease and the Aging Brain, Columbia University Medical Center, New York, NY, USA

**Keywords:** Haplotype Reference Panel (HRC), Imputation, Rare-Variant Aggregate Tests, Sequence data, Trans-Omics for Precision Medicine (TOPMed), UK Biobank

## Abstract

Imputation can cost-effectively generate genotypes for millions of variants. The power of performing rare variant aggregate association tests using imputed genotypes was evaluated. White Europeans from the UK Biobank with exome sequence and genotype array data were analyzed. Using the genotype type array data from the UK Biobank, imputation was performed using the HRC r1.1 (N=64,976 Haplotypes) and TOPMed r3 (N=267,194 haplotypes) reference panels. Simulations were used to compare the power of performing rare variant aggregate association analysis using sequence and imputed data. Additionally, the number of genes with ≥2 rare variants (missense, nonsense, splice site) was approximately the same for exome sequence and TOPMed imputed data, but HRC imputed data had ∼10% fewer genes. For the imputed data, using a less stringent R2 threshold (i.e., >0.3 vs. >0.8) led to greater power to detect aggregate associations due to additional rare variants included in the test. Exome sequence data provided the highest power for rare variant aggregate association testing, with TOPMed imputed variants usually having less than a 20% reduction in power. HRC imputed variants provided substantially less power. We also performed rare variant aggregate association analyses using UK Biobank phenotype, exome sequence data, and imputed variants for *PCSK9* and low-density lipoprotein (N=159,904 study subjects) and *APOC3* and triglyceride levels (N=160,036 study subjects). For these analyses, even when ultra-rare variants (minor allele frequency<0.001) were analyzed, significant aggregate associations could be detected for the exome sequence data and TOPMed and HRC imputed variants. Although there is a decrease in power for rare variant aggregate association tests when analyzing imputed variants compared to sequence data due to missing variants and uncertainty in genotypes, analysis of imputed data is a viable approach to detect rare variant aggregate associations when sequence data is unavailable.

## Introduction

Analysis of imputed data has become standard for conducting genome-wide association studies (GWAS). Performing imputation using genotype array data for a target sample and a reference imputation panel that consists of individuals with whole-genome sequence (WGS) data can provide genotypes to perform GWAS on tens of millions of variants. The additional variants obtained through imputation can increase the power of GWAS to detect associations and perform fine-mapping to identify causal susceptibility variants^1^.

Imputation can be performed using a variety of reference panels that include 1000 Genomes^2^, African Genome Resource^3^, Asthma among African-ancestry Populations in the Americas (CAAPA)^4^, Genome Asia Pilot (GAsP)^5^, HAPMAP2^6^, Haplotype Reference Consortium (HRC)^7^, Multi-ethnic HLA^8^, Southeast Asian Reference Database (SEAD)^9^, Trans-Omics for Precision Medicine (TOPMed)^10^, UK10K^11^, and the Westlake Biobank for Chinese (WBBC)^12^.

HRC and TOPMed imputation panels are two of the most frequently used. HRC (r1.1 2016) contains 32,470 individuals of predominately European ancestry and 39,635,008 variants while TOPMed (r3) has data on 133,597 reference samples from a variety of ancestry groups (∼55.1% European, ∼25.5% African/African American, ∼8.1% South Asian, ∼5.0% Hispanic/Latino, ∼3.5% East Asians, and ∼2.7% Middle Eastern) and 445,600,184 variants. Both HRC and TOPMed imputation panels contain WGS data for the autosomes and X chromosome. Imputation using a variety of reference panels can be performed on imputation servers, i.e. Michigan, NHLBI, Sanger, and Westlake, that allow users to upload their phased or unphased genotype data. These imputation servers, phase and impute each target sample separately, therefore the size of the target dataset does not impact imputation quality.

Imputation accuracy is dependent on both the imputation reference panel and the target sample. For the imputation reference panel sample size as well as ancestral diversity plays a role. For the target sample, marker density, quality of genotype data, as well as ancestry [linkage disequilibrium (LD) structure and representation on the imputation reference panel] also impact imputation performance^10^. It is also possible to perform meta-imputation^13^ where data from more than one reference panel is used to improve imputation accuracy and increase the number of variants. However, a caveat is that the reference panels must be available in the same build, which is not always the case, e.g., TOPMed is hg38 and HRC is hg37.

In addition to performing univariate analysis to detect associations, there is also an interest in performing aggregate analysis to detect rare-variant associations. Rare-variant aggregate association analysis has led to the discovery of genes involved in the etiology of lipid levels and coronary heart disease^14^ amongst others^15^. Currently, most rare-variant aggregate analysis has been limited to protein-coding regions due to limited availability of large samples with WGS data and uncertainty on how to aggregate rare variants in non-coding regions; though these challenges are starting to be addressed.^16^

Although some biobanks are generating exome sequence (ES) and/or WGS data (e.g., All of Us^17^ and UK Biobank^18^), many biobanks only have generated or plan to generate genotype array data (e.g., Finngen^19^, Estonian Biobank^20^, and Our Future Health) and have no immediate plans to generate either ES or WGS data for the entire sample. Additionally, for many genetic studies it is still cost-prohibitive to generate ES or WGS data for large sample sizes, i.e., hundreds of thousands of individuals. For many genetic studies and biobanks, genotype array data is available, or can readily be generated, allowing for the imputation of millions of variants.

Given the interest in performing rare-variant aggregate association tests for genetic studies, we investigated whether analyzing imputed data is a viable option. Unrelated White Europeans (N=168,206) from UK Biobank with ES, genotype array data, and imputed variants from the HRC (r1.1 2016) and TOPMed (Version r3) were used to investigate the feasibility of performing rare-variant aggregate analysis, using both simulation studies and analysis of low-density lipoproteins (LDL) and triglyceride (TG) levels. The distribution of rare variants and their quality (R^2^ - a measure of imputation accuracy and r^2^ – the correlation between genotypes obtained from imputation and sequence data) was also evaluated. Although the analysis of ES data was the most powerful approach, performing rare-variant [e.g. minor allele frequency (MAF) <0.001] aggregate association testing using imputed data is a viable alternative to analyzing sequence data.

## Materials and Methods

Two different arrays, the UK BiLEVE array (807,411 markers) and the UK Biobank Axiom array (825,927 markers), were used to assay samples from 49,950 and 438,427 UK Biobank^18^ study subjects, respectively. The intersection of the two arrays contains 733,322 autosomal variants. Study subjects missing ≥3% of their genotype data were removed. Variants with MAF>0.05 missing >5% of their genotypes and variants with MAF≤5% missing >1% of their genotypes were also removed. Self-reported ancestry and principal component (PC) analysis as previously described^21^ were used to obtain a sample of White Europeans. Using a kinship coefficient filter of 0.065 (3^rd^ degree relative or closer)^22^ related individuals within a relative set were removed so that unrelated individuals from a family could be retained. Variants that failed Hardy-Weinberg equilibrium (p<5×10^-15^) in the sample of unrelated White Europeans were also removed. A total of 651,867 variants remained after quality control (QC).

Using the Michigan and NHLBI imputation servers, imputation was performed, to generate variants with an R^2^>0.3, using the HRC^7^ (r1.1) and TOPMed^10^ (r3) reference panels, respectively. The target sample was phased using EAGLE2^23^ and imputation was performed for chromosomes 1, 2, and 11 using Minimac4^24^. To combine the imputed data from HRC and TOPMed, we retained the imputed variant with the highest R^2^ value.

UK Biobank ES data was generated using the IDT xGen Exome Research Panel v1.0 capture array, with different oligo lots used for the first (∼50K) and second release (∼150K) of exomes, that targeted 39Mbp of the human genome covering ∼19,400 genes, with coverage of >20x at 95.6% of the targeted bases^25,26^. The 200,643 samples from release 2 were processed using OQFE^25^ which aligned all the raw sequence data (FASTQs) to the human genome (GRCh38) using the Burrows-Wheeler Aligner (BWA)^27^ and Picard to mark duplicates. The Genome Analysis Toolkit (GATK)^28^ was used to call single nucleotide variants (SNVs) and insertion-deletions (indels), followed by base quality score recalibration (BQSR) to generate per-sample gVCFs. Multi-sample jointly called VCFs were generated using GLnexus^29^. Bcftools v.1.12^30^ was used to perform multiallelic splitting and left normalization. Variants with an allelic imbalance of <0.15 for SNVs and <0.2 for indels were removed. Genotypes with a read depth <10x or a genotype quality score <20 were also removed. Additionally, those variants that were missing >10% of their genotypes were removed. Gene regions as well as stop-gain, stop-loss, start-loss, start-gain, missense, splice site, and frameshift variants were defined using ANNOVAR^31^ and extracted from each dataset, (i.e., ES and HRC and TOPMed imputed data). The variants were annotated with allele frequencies obtained from gnomAD^32^ for non-Finnish Europeans (NFE). Missense and splice site variants were also annotated with CADD c-scores^33^.

We evaluated rare loss of function (LoF) variants (stop-gain, stop-loss, start-loss, start-gain, frameshift) and missense and splice sites with and without a CADD c-score >20. Variants obtained from ES, HRC imputed, TOPMed imputed data as well as combined panels of variants, i.e., HRC_TOPMed and ES_HRC_TOPMed were evaluated. Variants with the highest R^2^ values regardless of the imputation panel used were included in the HRC_TOPMed sample. For the ES_HRC_TOPMed sample, missing ES variants were replaced with either an HRC or TOPMed imputed variant depending on which one had the highest R^2^ value.

For chromosomes 1 and 2, the rare variant allele frequency distribution by variant type was compared for ES and imputed variants, where variant frequencies were obtained from gnomAD NFE using five thresholds, i.e. MAF≤1×10^-5^; (1×10^-5^<MAF ≤1×10^-4^); (1×10^-4^<MAF≤1×10^-3^); (1×10^-3^< MAF≤5×10^-3^); and (5×10^-3^< MAF<1×10^-2^). For rare-variant aggregate tests gnomAD NFE frequencies were used to determine rare variant thresholds, i.e., MAF<0.001, <0.005, and <0.01 to establish which rare variants within a gene region should be included in the aggregate test. For the imputed data, two R^2^ thresholds were examined, i.e., >0.3 and >0.8. We assessed the number of genes with ≥2 rare variants in both the ES and imputed data. And we examined the R^2^ and r^2^ values and their standard errors for HRC and TOPMed imputed variants. A two-sided Welch two-sample T-test was used to determine if there was a statistical difference between mean (μ) R^2^ values obtained from HRC and TOPMed and between μ R^2^ and r^2^ values obtained from the same dataset.

For the ES and imputed data, type 1 error was evaluated at three different MAF thresholds and for imputed variants the two R^2^ thresholds. To evaluate type I error, from the 168,206 unrelated White Europeans we randomly assigned 40,000 individuals as cases and 60,000 as controls for each gene. We tested every gene on chromosomes 1 and 2 with ≥2 rare variants (LoF, missense and splice site) using the three rare variant MAF thresholds and two R^2^ thresholds for imputed variants. Rare-variant aggregate analysis was performed using logistic regression by applying the Burden of Rare Variants (BRV)^34^ test, which is a fixed effects test that regresses the number of rare variants observed for each sample on the outcome of interest. For imputed variants, the dosages were summed for every variant within each gene region for every individual.

To evaluate power, we again generated samples of 40,000 cases and 60,000 controls for every gene on chromosome 1 and 2 with ≥2 rare variants, this time using disease prevalences of 0.1 and 0.2, a variety of odds ratios (ORs) i.e., 1.2., 1.5, and 1.8, and the proportion of variants selected within a gene region to be causal, i.e., 1.0, 0.75, and 0.50. We performed analysis for the three rare variant MAF and two R^2^ thresholds using LoF variants, missense and splice site. All genes with ≥2 variants were analyzed using the same logistic regression framework as was used to evaluate type I error. Power was estimated as the number genes that met an exome-wide significance level of 2.5×10^-6^ divided by the maximum number of genes tested in all datasets, i.e., ES, HRC, TOPMed, HRC_TOPMed, and ES_HRC_TOPMed. For those genes that met the exome-wide significance, we also evaluated the μ R^2^ per gene for the HRC and TOPMed imputed variants. To evaluate whether the loss in power of imputed data compared to ES data was due to genotype uncertainty or missing data, we considered only those variants that overlapped between ES and TOPMed imputed data for the three MAF thresholds and R^2^>0.3.

Rare-variant aggregate associations between LDL levels (field 30780) and *PCSK9* (chromosome 1p32.3) and TG levels (field 30870) and *APOC3* (chromosome 11p23.3) were tested. After removing 6,541 individuals who used statins, 1,761 individuals with missing LDL levels, and 1,629 individuals with missing TG levels, there were 159,904 and 160,036 study subjects to analyze with LDL and TG levels, respectively. Association analysis of the log-transformed LDL and TG levels were performed using linear regression applying the rare-variant aggregate fixed effect BRV test, analyzing the number of variants or the sum of the dosages within a gene region for ES and imputed data, respectively. Covariates in the linear regression model included sex, age at time of the LDL or TG measurement, and two PCs. The PCs were generated^35^ using the pruned genotype array data (r^2^<0.2 and MAF>0.01). ES and imputed variants were analyzed using three rare variant MAF thresholds as well as the two R^2^ thresholds for imputed variants. Variant allele frequencies for inclusion in the rare-variant aggregate association tests were based on gnomAD NFE. Rare-variant aggregate association analysis was performed analyzing LoF, missense, and splice variants as well as LoF variants and missense and splice site variants with a CADD c-score >20.

## Results

### Rare variants in exome sequence and imputed data

Rare-variant aggregate association analysis is usually performed on regions with at least two variants; therefore, we compared the number of genes on chromosomes 1 and 2 that met this criterion (Table 1). The number of genes with at least two LoF, missense, or splice site variants for a MAF < 0.01 was the highest for imputed TOPMed variants (R^2^>0.3, N=3,186 genes) with ES having slightly fewer genes (N=3,146), but the number of genes decreased for TOPMed if the R^2^ threshold was increased to 0.8 (N=3,102 genes). The fewest number of genes (N=2,271) with ≥2 variants was observed for HRC (R^2^>0.8). The number of genes with ≥2 rare variants (MAF <0.001) when a CADD c-score criterion was applied, (i.e., >20 for missense and splice variants) fell slightly for ES (0.13%) and TOPMed (R^2^>0.3, 0.57%), however for HRC the decrease in the number of genes was substantially greater (R^2^>0.3, 10.18%) (Table 1).

**Table 1.** Number of genes with at least two rare variants.

| Dataset | R<sup>2</sup> | MAF < 0.01 |  | MAF < 0.005 |  | MAF < 0.001 |  |
| --- | --- | --- | --- | --- | --- | --- | --- |
|  |  | LoF, missense & splice site | LoF, (missense & splice site with CADD≥20) | LoF, missense & splice site | LoF, (missense & splice site with CADD≥20) | LoF, missense & splice site | LoF, (missense & splice site with CADD≥20) |
| ES | - | 3,146 | 3,142 | 3,146 | 3,142 | 3,146 | 3,142 |
| HRC | >0.3 | 2,922 | 2,656 | 2,905 | 2,637 | 2,819 | 2,532 |
|  | >0.8 | 2,271 | 1,896 | 2,188 | 1,816 | 1,819 | 1,414 |
| TOPMed | >0.3 | 3,186 | 3,169 | 3,186 | 3,169 | 3,185 | 3,167 |
|  | >0.8 | 3,102 | 3,043 | 3,099 | 3,040 | 3,091 | 3,025 |
| HRC_TOPMed | >0.3 | 3,198 | 3,177 | 3,198 | 3,177 | 3,196 | 3,174 |
|  | >0.8 | 3,108 | 3,049 | 3,105 | 3,046 | 3,097 | 3,033 |
| ES_HRC_TOPMed | >0.3 | 3,203 | 3,176 | 3,203 | 3,176 | 3,203 | 3,175 |
|  | >0.8 | 3,181 | 3,163 | 3,181 | 3,163 | 3,180 | 3,163 |

Displayed for chromosomes 1 and 2 are the number of genes with $\geq 2$ variants with MAF below the rare variant threshold of 0.01, 0.005, and 0.001. The MAF for each variant is obtained from gnomAD for NFE. **Abbreviations:** CADD, combined annotation–dependent depletion; ES, exome sequence; gnomAD, Genome Aggregation Database; HRC, Haplotype Reference Consortium; LoF, loss of function; MAF, minor allele frequency; NFE, non-Finish Europeans; R<sup>2</sup>, imputation quality; and TOPMed, Trans-Omics for Precision Medicine.

The number of variants (LoF, missense, and splice site for chromosomes 1 and 2) for the imputed data varied greatly depending on the imputation reference panel and the R^2^ threshold (Tables 2 and S1, Figures 1 and S1). For example, when the R^2^ threshold for the HRC imputed data was increased from 0.3 to 0.8, the number of variants decreased by 57.3%. This shift towards higher R^2^ thresholds also resulted in the exclusion of rarer variants, especially for variants with MAF<1×10^-4^ (Figures 1 and S1, Table 2 and S1). The μ MAF for ES data is smaller than that for the imputed datasets, e.g., for instance, for MAF <0.001, the ES data had 816,399 variants (μ MAF=3.73×10^-5^), whereas HRC and TOPMed (R^2^>0.3) had 28,674 variants (μ MAF=2.17×10^-4^) and 235,264 variants (μ MAF=6.09×10^-5^), respectively. Both ES data and TOPMed imputed dataset contain a large proportion of ultra-rare variants with MAF<1×10^-5^ compared to HRC (Figures 1 and S1, Tables 2 and S1).

**Figure 1.**
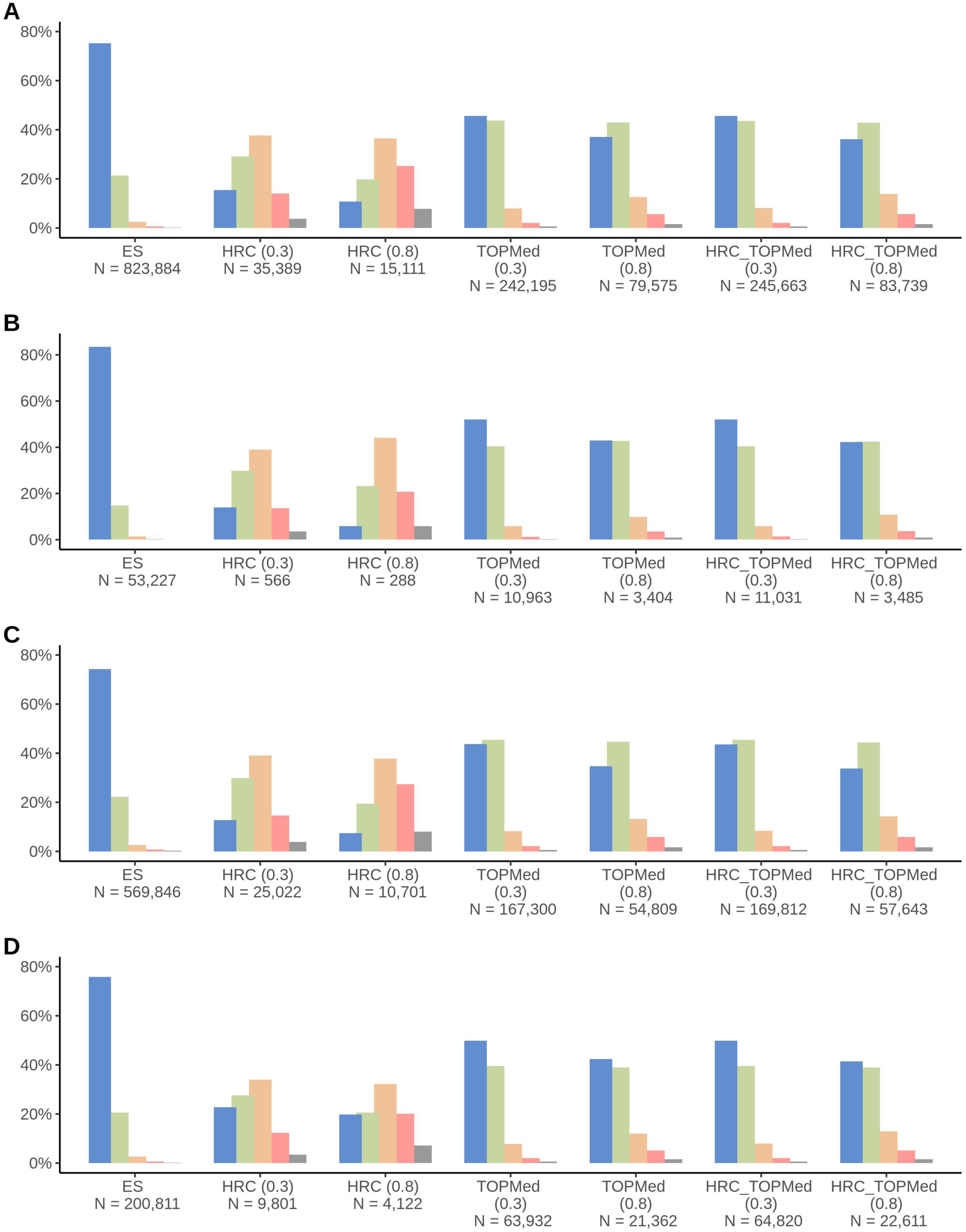
Bar chart displaying the distribution of variant minor allele frequencies. The minor allele frequency (MAF) categories are shown in the following colors: ***Blue*** (MAF ≤ 1×10^-5^; ***Green*** (1×10^-5^<MAF≤1×10^-4^); ***Orange*** (1×10^-4^<MAF≤1×10^-3^); ***Pink*** (1×10^-3^< MAF≤5×10^-3^); and ***Grey*** (5×10^-3^<MAF<1×10^-2^). The numbers in brackets represent the R^2^ threshold and N is the number of variants in each MAF category. **Abbreviations**: ES, exome sequence; HRC, Haplotype Reference Consortium; LoF, loss of function; and TOPMed, Trans-Omics for Precision Medicine. For details on the percentage of variants in each MAF category, refer to Table S8. In the panels are displayed the distribution by variant type: **(A)** all (LoF, missense, and splice site); **(B)** LoF; **(C)** missense; and **(D)** splice site.

**Table 2.**
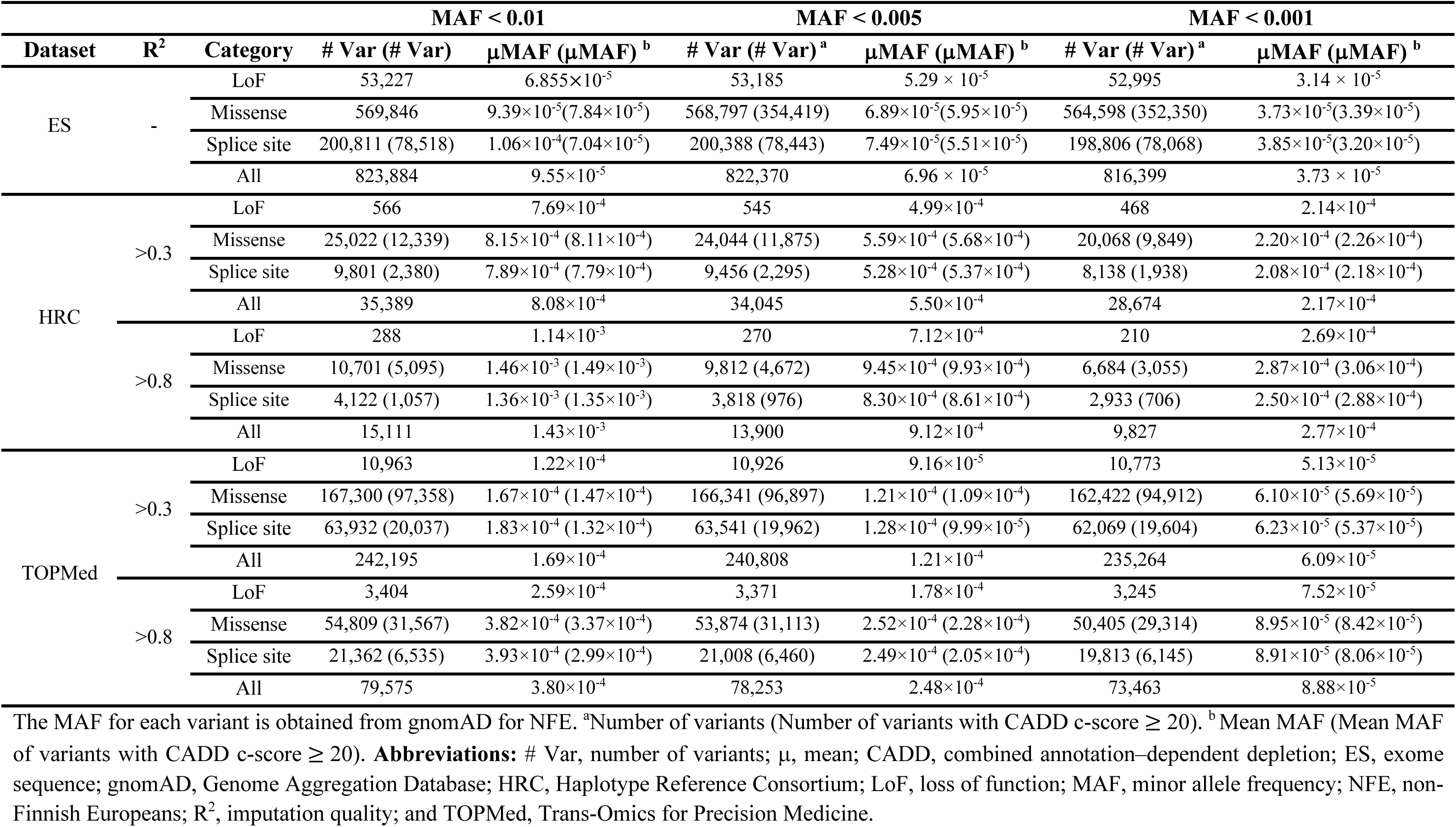
Number of variants for each rare variant minor allele frequency threshold and their mean minor allele frequencies.

The largest proportion of uniquely shared variants (LoF, missense, and splice site) can be found between the ES and TOPMed (R^2^>0.3, N=161,571) datasets (Figure S2). Conversely, only a fraction of variants was uniquely shared between ES and HRC data (R^2^>0.3, N=2,421). The ES data had 629,566 unique variants, while TOPMed and HRC (R^2^>0.3) had 48,803 and 1,047 unique variants, respectively (Figure S2).

The R^2^ values were lowest for the variants with MAF<0.001 for HRC, TOPMed, and HRC_TOPMed. There are 8.2X more TOPMed than HRC imputed variants with an MAF<0.001. The large difference in the number of variants observed in TOPMed compared to HRC is that TOPMed has a greater preponderance of variants with an MAF<0.001. Due to the larger number of rare variants (MAF<0.001) in TOPMed the μ R^2^ values (0.667) are lower than observed for HRC (0.689) (p-value=6.46×10^-67^). However, for imputed variants with MAF<0.001 that overlapped between HRC and TOPMed, TOPMed (μ R^2^=0.734) imputation accuracy was higher than for HRC (μ R^2^=0.706) (p-value=1.87×10^-63^) (Tables 3 and S2). For the HRC_TOPMed imputed variants (MAF<0.001) the number of variants and μ R^2^ values were higher than for variants only imputed using TOPMed (Table 3). Regardless of the imputation panel used there were <6,000 variants with a 0.001≤MAF<0.005 and <1,500 variants with a 0.005≤MAF<0.01, for these categories the μ R^2^ was always >0.88 with TOPMed and HRC (Table 3 and S2). For almost all categories the μ R^2^ and r^2^ values were significantly different (Table S3), but one was not consistently higher than the other (Table 3).

**Table 3.** Mean R^2^ and r^2^ for HRC and TOPMed variants.

| Dataset | MAF | $R^2$ | | $r^2$ | |
| --- | --- | --- | --- | --- | --- |
| | | $\mu$ (SE) | # Var | $\mu$ (SE) | # Var |
| HRC | MAF < 0.001 | 0.689 (0.001) | 28,674 | 0.669 (0.002) | 26,285 |
|  | MAF < 0.001<sup>a</sup> | 0.706 (0.001) | 25,438 | 0.690 (0.002) | 24,029 |
|  | 0.001 ≤ MAF < 0.005 | 0.882 (0.002) | 5,371 | 0.905 (0.002) | 5,256 |
|  | 0.001 ≤ MAF < 0.005<sup>a</sup> | 0.889 (0.002) | 5,190 | 0.913 (0.002) | 5,127 |
|  | 0.005 ≤ MAF < 0.01 | 0.930 (0.003) | 1,344 | 0.938 (0.006) | 1,306 |
|  | 0.005 ≤ MAF < 0.01<sup>a</sup> | 0.939 (0.002) | 1,293 | 0.943 (0.005) | 1,270 |
| TOPMed | MAF < 0.001 | 0.667 (0.000) | 235,264 | 0.653 (0.001) | 185,099 |
|  | MAF < 0.001<sup>a</sup> | 0.734 (0.001) | 25,438 | 0.833 (0.001) | 24,029 |
|  | 0.001 ≤ MAF < 0.005 | 0.903 (0.001) | 5,544 | 0.936 (0.001) | 5,447 |
|  | 0.001 ≤ MAF < 0.005<sup>a</sup> | 0.910 (0.001) | 5,190 | 0.945 (0.001) | 5,127 |
|  | 0.005 ≤ MAF < 0.01 | 0.950 (0.002) | 1,387 | 0.948 (0.006) | 1,351 |
|  | 0.005 ≤ MAF < 0.01<sup>a</sup> | 0.958 (0.002) | 1,293 | 0.959 (0.005) | 1,270 |
| HRC_TOPMed | MAF < 0.001 | 0.673 (0.000) | 238,500 | 0.648 (0.001) | 187,355 |
|  | 0.001 ≤ MAF < 0.005 | 0.921 (0.002) | 5,725 | 0.937 (0.002) | 5,576 |
|  | 0.005 ≤ MAF < 0.01 | 0.946 (0.003) | 1,438 | 0.943 (0.006) | 1,387 |

Displayed for imputed LoF, missense, and splice site variants from chromosomes 1 and 2. The MAF for each variant is obtained from gnomAD for NFE. Only variants with a $R^2 > 0.3$ are included. The $r^2$ values can only be calculated for variants that were included in the exome sequence data. <sup>a</sup> Only imputed variants that overlap between HRC and TOPMed are included. **Abbreviations:** # Var, number of variants; $\mu$ , mean; gnomAD, Genome Aggregation Database; HRC, Haplotype Reference Consortium; MAF, minor allele frequency; NFE, non-Finnish European; $r^2$ , correlation between imputed and exome sequence data variants; $R^2$ , imputation quality; SE, standard error, and TOPMed, Trans-Omics for Precision Medicine.

### Type I error

Data was generated under the null to evaluate type I error. No deflation of p-values was observed, and type I errors were well controlled (Figure S3).

### Power to detect rare-variant aggregate associations

Simulation studies were performed to evaluate the power of rare-variant aggregate association test using a variety of parameters (Tables S4 and S5, Figures 2 and S4). The percent change in power for the imputed data (HRC and TOPMed) was also compared to the ES data (Tables S6 and S7). The power when analyzing imputed data (HRC and TOPMed) plus ES data was slightly greater than analyzing ES data alone (Table S4-S7, Figures 2 and S4). There was a drop in power when only imputed data were analyzed, with HRC_TOPMed imputed data having slightly greater power than analyzing TOPMed imputed data alone. For example, for a disease prevalence of 0.1 and an OR of 1.5 when all rare variants (MAF<0.01) within a gene region are causal the power decreased for HRC_TOPMed (R^2^>0.3) imputed data compared to ES data by 13.61% and for TOPMed imputed data compared to ES data by 15.44% (Tables S6). Lower power was obtained from analyzing HRC imputed data (Table S4-S7, Figure 2 and S4) compared to TOPMed imputed data. In all cases analyzing imputed data with an R^2^>0.3 provided greater power than using an R^2^ threshold of 0.8 (Tables S4-S7, Figures 2 and S4). When the MAF threshold was reduced from 0.01 to 0.005 there was a decrease in power for all data types. For example, for a disease prevalence of 10%, and rare variants with an OR=1.5, i.e., the power reduction for ES was 5.73%, TOPMed (R^2^>0.3) 10.41%, and HRC (R^2^>0.3) 18.87%. The power reduction was much greater for the same scenario when the MAF threshold was reduced from 0.005 to 0.0001, i.e. ES (21.33%), TOPMed (R^2^>0.3; 39.48%), and HRC (R^2^>0.3; 59.63%) (Table S4 and Figure S4). This loss of power was due to variants being removed when the MAF threshold was reduced from 0.005 to 0.001, ES (0.70%), TOPMed (2.10%), and HRC (14.00%) (Tables S8 and S9). However, for imputed variants, the loss in power was not only due to fewer variants being available for analysis but also the decrease on the μ R^2^ values as the MAF thresholds decreased (Table S10). We also compared power obtained when performing rare-variant aggregate association analysis only analyzing variants that overlap between ES and TOPMed imputed data as well as all available ES variants. There was a slightly greater loss in power due to missing variants than due to analyzing dosages, e.g., for an OR=1.5, disease prevalence of 0.1, all variants are causal, and a rare variant MAF threshold of 0.005 was applied there was a 21.2% drop in power when only analyzing the ES variants that were also available in TOPMed (R^2^>0.3) imputed data compared to analyzing all variants which were available in the ES data. There was an additional 11.4% loss in power when analyzing these same variants but analyzing the dosages obtained from imputation (Table S11).

**Figure 2.**
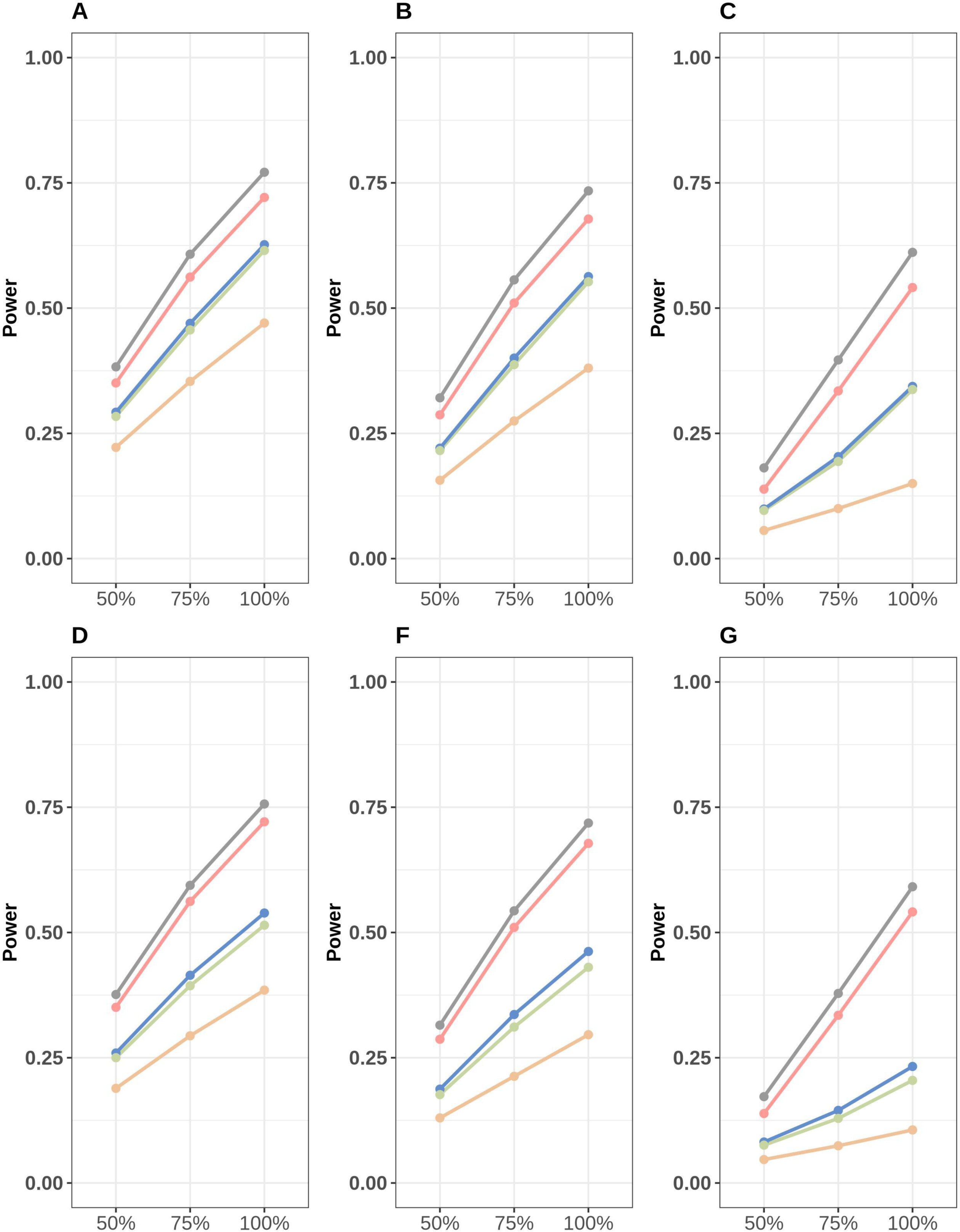
Power to detect rare-variant aggregate associations for exome sequence and imputed variant data. Power was evaluated by generating cases (N=40,000) and controls (N=60,000) for each gene on chromosomes 1 and 2 with at least two variants. The disease prevalence was 0.2, the proportion of causal rare variants with each gene region varied between 100% and 50%, with each causal rare variant within a gene region having an odds ratio of 1.5. Power was estimated as the number genes that met an exome-wide significance level of 2.5×10^-6^ divided by the maximum number of genes tested for ES, HRC, TOPMED, HRC_TOPMed, or ES_HRC_TOPMed. The data type is represented by the following colors: ***Pink*** (ES); ***Orange*** (HRC); ***Green*** (TOPMed); ***Blue*** (HRC_TOPMed); ***Grey*** (ES_HRC_TOPMed). **Abbreviations**: ES, exome sequence; HRC, Haplotype Reference Consortium; MAF, minor allele frequency; and TOPMed, Trans-Omics for Precision Medicine. The R^2^ value for imputed variant for panels: **A-C** R^2^>0.3 and **D-F** R^2^>0.8. The following rare variant MAF thresholds were used for the analysis: MAF<0.01 panels **A** and **D**; MAF<0.005 panels **B** and **E**; and MAF<0.001 panels **C** and **F**.

### LDL and TG level rare-variant aggregate associations with PCSK9 and APOC3

For both the rare variant aggregate analyses of *PCSK9* and *APOC3,* regardless of the data type, the same results were obtained when a rare variant MAF threshold of <0.01 or <0.005 was used since the reduction in MAF did not result in any additional variants being removed. The most significant results were obtained when either ES data or ES_HRC_TOPMed imputed variants were analyzed, limiting the analysis to LoF variants, and missense, and splice site variants with a CADD c-score ≥20, i.e., *APOC3* association with TG levels p-value=1.05×10^-262^ and *PCSK9* association with LDL levels p-value=8.25×10^-93^. For aggregate analysis of imputed variant for this same MAF threshold the most significant results for TOPMed (R^2^>0.3) when LoF variants, and missense, and splice site variants with a CADD c-score ≥20 were analyzed, i.e., *APOC3* association with TG levels p-value=4.04×10^-197^ and *PCSK9* association with LDL levels p-value=1.29×10^-54^, while for HRC the *APOC3* association with TG levels p-value=3.38×10^-182^ and *PCSK9* association with LDL levels p-value=1.91×10^-12^. For all analyses increasing the R^2^ threshold to >0.8 led to less significant results for both TOPMed and HRC. The p-values for both *APOC3* and *PCSK9* increased when the MAF threshold was reduced to 0.001, due to a strongly associated variant being removed. For example, for ES data when LoF and missense and splices variants with a CADD c-score ≥20 were analyzed the number of variants in *APOC3* dropped from 26 to 25 variants, and the p-value for an association with TG levels increased to 9.62×10^-48^. For the same analysis but using TOPMed (R^2^>0.3) imputed variants the number of *APOC3* variants decreased from 10 to 9 and the p-value increased to 2.25×10^-28^, while for HRC (R^2^>0.3) the number of variants decreased from 3 to 2 and the p-value increased to 3.39×10^-24^(Table 4). The *APOC3* splice site variant (11:116701354 IVS2+1G→A) which was removed due to its MAF being >0.001 was tested and significant associations were observed for ES (p-value=2.48×10^-219^), HRC (R^2^=0.836; p-value=1.17×10^-161^), and TOPMed (R^2^=0.854; p-value=4.47×10^-175^) with TG levels. This splice site variant has previously been reported to reduce TG levels^36^ and has an MAF=2.3×10^-3^ for NFE in gnomAD.

**Table 4.** Rare variant aggregate association analysis of triglyceride levels and *APOC3* and low-density lipoprotein levels and *PCSK9*.

| Dataset | R<sup>2</sup> | APOC3 |  |  |  | PCSK9 |  |  |  |
| --- | --- | --- | --- | --- | --- | --- | --- | --- | --- |
|  |  | LoF, missense & splice site |  | LoF, (missense & splice site with CADD≥20) |  | LoF, missense & splice site |  | LoF, (missense & splice site with CADD≥20) |  |
|  |  | # Var | p-value | # Var | p-value | # Var | p-value | # Var | p-value |
| MAF < 0.01 and MAF < 0.005 <sup>a</sup> |  |  |  |  |  |  |  |  |  |
| ES | - | 60 | 3.07×10<sup>-234</sup> | 26 | 1.05×10<sup>-262</sup> | 394 | 5.47×10<sup>-71</sup> | 200 | 8.25×10<sup>-93</sup> |
| HRC | >0.3 | 3 | 3.38×10<sup>-182</sup> | 3 | 3.38×10<sup>-182</sup> | 16 | 1.94×10<sup>-12</sup> | 6 | 1.91×10<sup>-12</sup> |
|  | >0.8 | 2 | 9.97×10<sup>-170</sup> | 2 | 9.97×10<sup>-170</sup> | 6 | 6.99×10<sup>-11</sup> | 2 | 4.56×10<sup>-15</sup> |
| TOPMed | >0.3 | 18 | 1.34×10<sup>-190</sup> | 10 | 4.04×10<sup>-197</sup> | 124 | 2.40×10<sup>-38</sup> | 61 | 1.29×10<sup>-54</sup> |
|  | >0.8 | 4 | 1.76×10<sup>-183</sup> | 4 | 1.76×10<sup>-183</sup> | 34 | 7.40×10<sup>-30</sup> | 15 | 1.23×10<sup>-44</sup> |
| HRC_TOPMed | >0.3 | 18 | 4.15×10<sup>-187</sup> | 10 | 4.60×10<sup>-194</sup> | 125 | 1.15×10<sup>-39</sup> | 61 | 3.75×10<sup>-54</sup> |
|  | >0.8 | 4 | 6.41×10<sup>-183</sup> | 4 | 6.41×10<sup>-183</sup> | 38 | 5.15×10<sup>-32</sup> | 16 | 3.68×10<sup>-45</sup> |
| ES_HRC_TOPMed | >0.3 | 65 | 3.67×10<sup>-231</sup> | 28 | 8.54×10<sup>-260</sup> | 412 | 8.87×10<sup>-71</sup> | 204 | 2.55×10<sup>-93</sup> |
|  | >0.8 | 60 | 3.07×10<sup>-234</sup> | 26 | 1.05×10<sup>-262</sup> | 394 | 5.47×10<sup>-71</sup> | 200 | 8.25×10<sup>-93</sup> |
| MAF < 0.001 |  |  |  |  |  |  |  |  |  |
| ES | - | 59 | 5.47×10<sup>-32</sup> | 25 | 9.62×10<sup>-48</sup> | 393 | 9.35×10<sup>-53</sup> | 199 | 2.53×10<sup>-74</sup> |
| HRC | >0.3 | 2 | 3.39×10<sup>-24</sup> | 2 | 3.39×10<sup>-24</sup> | 15 | 2.39×10<sup>-9</sup> | 5 | 5.73×10<sup>-12</sup> |
|  | >0.8 | 1 | NA | 1 | NA | 6 | 6.99×10<sup>-11</sup> | 2 | 4.56×10<sup>-15</sup> |
| TOPMed | >0.3 | 17 | 1.39×10<sup>-25</sup> | 9 | 2.25×10<sup>-28</sup> | 123 | 5.23×10<sup>-26</sup> | 60 | 9.89×10<sup>-42</sup> |
|  | >0.8 | 3 | 1.49×10<sup>-13</sup> | 3 | 1.49×10<sup>-13</sup> | 33 | 5.72×10<sup>-17</sup> | 14 | 3.98×10<sup>-34</sup> |
| HRC_TOPMed | >0.3 | 17 | 2.63×10<sup>-20</sup> | 9 | 2.23×10<sup>-23</sup> | 124 | 4.19×10<sup>-27</sup> | 60 | 3.14×10<sup>-41</sup> |
|  | >0.8 | 3 | 1.49×10<sup>-13</sup> | 3 | 1.49×10<sup>-13</sup> | 37 | 3.53×10<sup>-19</sup> | 15 | 1.15×10<sup>-34</sup> |
| ES_HRC_TOPMed | >0.3 | 64 | 1.19×10<sup>-30</sup> | 27 | 1.48×10<sup>-45</sup> | 411 | 1.37×10<sup>-52</sup> | 203 | 7.68×10<sup>-75</sup> |
|  | >0.8 | 59 | 5.47×10<sup>-32</sup> | 25 | 9.62×10<sup>-48</sup> | 393 | 9.35×10<sup>-53</sup> | 199 | 2.53×10<sup>-74</sup> |

<sup>a</sup> MAF thresholds < 0.01 and < 0.005 provide identical results. **Abbreviations:** # Var: number of variants; *APOC3*: Apolipoprotein C-III; CADD: combined annotation–dependent depletion; ES: exome sequence; HRC: Haplotype Reference Consortium; LoF: loss of function; MAF: minor allele frequency; *PCSK9*: Proprotein Convertase Subtilisin/Kexin Type 9; $R^2$ threshold: imputation quality; and TOPMed: Trans-Omics for Precision Medicine.

## Discussion

Our findings demonstrate that using a threshold of R^2^ >0.3 usually provided greater power than an R^2^>0.8 without increasing type I error. An R^2^>0.8 led to the exclusion of variants, particularly those in the lower frequency range, due to these variants being less likely to be imputed with high accuracy (Figure 1). Therefore, we suggest using a threshold of R^2^>0.3 for rare-variant aggregate association analysis. Power to detect rare-variant aggregate associations was higher using TOPMed than HRC, combining the imputed variants generated from both imputation panels lead to a slight increase in power compared to only analyzing imputed variant obtained from the TOPMed panel. Although there was a loss of power when analyzing imputed variants compared to ES data, the number of gene regions that could be tested, was approximately the same for the ES and TOPMed imputed data, with slightly more gene regions for TOPMed (R^2^>0.3) imputed data, i.e., 1.26%. Although there was a power loss analyzing imputed rare variants with adequately large sample sizes, this is a powerful approach to detect associations.

Rare-variant aggregate studies use a variety of MAF thresholds, with the upper bound ranging from 0.01 to 0.001. For larger datasets, such as the UK Biobank, it is often possible to detect associations with individual variants that are on the rare spectrum, e.g., 0.001<MAF< 0.01 even if their effect size is modest, e.g., OR=1.2. For rare-variant aggregate tests, it is beneficial to test variants in aggregate that due to either their MAF or in some cases their effect sizes, associations cannot be detected using a single variant association test. Therefore, with sufficiently large samples a MAF <0.001 can be used for rare-variant aggregate association tests.

Analyzing imputed variant data is less powerful than directly analyzing sequence data due to two factors: 1) Variants are missing from the imputed data, particularly ultra-rare variants, e.g., MAF<5×10^-5^; and 2) True genotypes of the imputed variants are unknown, instead only probabilities for the three possible genotypes are available. These probabilities are usually used to obtain dosages which are analyzed. With decreasing imputation accuracy, e.g., R^2^ the informativeness of imputed variants decreases compared to a variant obtained from sequencing (assuming there is no genotyping error). For imputed data the μ R^2^ decreases with decreasing MAF. For TOPMed compared to HRC, not only was it possible to impute more variants but their R^2^ values were also higher.

When analyzing ES data, due to some regions not being included in the capture array or removal of variants due to QC, power can be increased by substituting missing variants in ES data with imputed genotypes. However, the benefits of including imputed variants when analyzing ES data will depend on the size of the exome capture array and read depth, with the increase in power diminishing when ES data is generated using higher read depths and larger capture arrays.

Unlike for sequence data, imputed variants on the genotype level do not have missing data. Missing genotypes can reduce power to detect associations, and for rare-variant aggregate tests if the percent of missing data is greater in either cases or controls type I error can be increased^34^. We replaced missing genotypes in the ES data with imputed variants, however due to low rates of missing genotypes, the impact on power was negligible (data not shown).

For the rare-variant aggregate testing, we only used a fixed effect test, i.e. BRV. We did not use random effects tests, e.g., SKAT^37^ or the omnibus SKAT-O test^38^. Our goal was to compare analyzing sequence data to imputed data and not to compare rare-variant aggregate methods. Power will be impacted for all rare-variant aggregate tests by variant effect sizes, number and frequencies of the variants, and the proportion of benign variants.

Although UK Biobank has imputed data available, we re-imputed the data for several reasons. The UK Biobank performed imputation using the TOPMed version r2 reference panel that has ∼36,300 fewer individuals than the current TOPMed version r3 reference panel. The UK Biobank also performed imputation using a reference panel consisting of HRC, UK10K, and 1,000 genomes, performing imputation using Impute2^39^. Since this reference panel is not publicly available, we performed imputation just using only HRC. Additionally, for consistency we generated both the HRC and TOPMed imputed data using Minimac4.

This study was limited to evaluating rare imputed variant aggregate association testing for a European population. The ability to impute variants for non-European populations, i.e. African Americans, Asians (East and South), and Admixed Americans has already been evaluated for the TOPMed (version r2) and HRC imputation reference panels. HRC imputed rare variants poorly for all non-European populations. For TOPMed, the ability to impute rare variants was comparable for African Americans, Admixed Americans, and Europeans, with the imputation accuracy being much lower for Asians (East and South). HRC has very low representation on non-European populations and although TOPMed (r2) imputation panel did include Asians their numbers were low, most likely impacting these panels’ ability to impute variants for non-Europeans (HRC) and Asians (TOPMed r2)^10^.

It would be advantageous to have larger imputation reference panels with individuals of diverse ancestry to improve imputation accuracy. In many cases these resources of WGS data to generate imputation panels are already available or will be generated in the next few years. TOPMed currently has WGS data on ∼180k individuals and has a targeted goal of sequencing 220k ancestry-diverse samples, which could be used to increase the size of the current imputation panel (N=133,597). GAsP (N=1031) plans to generate an imputation panel using 100,000 samples of diverse Asian ancestry. All of Us will generate WGS data on at least 1,000,000 study subjects of diverse ancestry and currently there are 242,580 WGS samples (e.g., N=50,080 African American 41,940 Hispanic; 7,440 Asian) available for analysis. Additionally, the UK Biobank, which is predominantly White European has generated WGS data on ∼500k individuals. Even if these large samples of WGS data become available for imputation, there are still populations without sufficient representation e.g., East Africans and Indigenous South American populations.

The analysis of imputed variants can be used as a post-analysis QC measure. If analysis of genotype or sequence data yields a statistically significant association, but with no other associated variants in the region, this could signify that the observed signal occurred due to genotyping error. Within small genomic regions, due to LD structure, there are usually clusters of associated variants. For variants with genotype errors there often is greatly reduced LD with nearby variants, thus leading to the observation of only a single variant being associated. For this situation, imputed data for the variant can be analyzed to determine if there is concordance between results (with genotype or sequence data), when results greatly differ this could signify that the observed association is a false positive. When performing rare-variant aggregate tests using either WGS or ES data if the signal is not driven by ultra-rare variants, which could not be imputed, rare-variant aggregate association analysis can be performed using imputed variants, instead of the variants obtained from sequence data, to assess if the signal remains.

In conclusion, when sequence data is unavailable, the analysis of imputed data is a cost-effective approach to detect rare-variant aggregate associations. When performing rare-variant aggregate association testing analyzing variants imputed using more than one imputation panel can increase power. As larger diverse imputation panels become available, the power to detect rare-variant aggregate associations will increase, particularly for populations that are not well represented on currently available imputation panels.

## Supporting information

https://github.com/Suzannemleal/supplemental-material.git

## Web Resources

ANNOVAR: https://annovar.openbioinformatics.org/en/latest/

Combined Annotation Dependent Depletion (CADD): https://cadd.gs.washington.edu/

Genome Aggregation Database (gnomAD): https://gnomad.broadinstitute.org/

Haplotype Reference Consortium: https://www.sanger.ac.uk/collaboration/haplotype-reference-consortium/

Michigan Imputation Server: https://imputationserver.sph.umich.edu/index.html#!

NHLBI Trans-Omics for Precision Medicine (TOPMed): https://topmed.nhlbi.nih.gov/

Sanger Imputation Service: https://www.sanger.ac.uk/tool/sanger-imputation-service/

TOPMed Imputation Server: https://imputation.biodatacatalyst.nhlbi.nih.gov/#!

UK Biobank: https://www.ukbiobank.ac.uk/

Westlake Imputation Server: https://imputationserver.westlake.edu.cn/

## Acknowledgments

This research was conducted using data from UK Biobank (project IDs 19746 and 32285), a major biomedical database, under generic approval from the National Health Services’ National Research Ethics Service. UK Biobank is generously supported by its founding funders the Wellcome Trust and UK Medical Research Council, as well as the Department of Health, Scottish Government, the Northwest Regional Development Agency, British Heart Foundation, and Cancer Research UK.

This work was supported by a grant from the National Institute of Deafness and Other Communications Disorders (NIDCD) R01DC017712 to S.M.L, P.L.A., and A.T.D.

## Supplemental data

Four supplemental figures and 11 supplemental tables

## Data and code availability

UK Biobank exome sequence and genotype data used for this study are available to approved researchers. Instructions for access to UK Biobank data are available at https://www.ukbiobank.ac.uk/enable-your-research. The code used to process the data and perform the analysis are available at https://github.com/statgenetics/imputation

