## Supplementary material for "Rare-variant aggregate association analysis using imputed data is a powerful approach": https://github.com/Suzannemleal/supplemental-material.git

**Supplementary Figure 1.** Bar chart displaying the distribution of missense and splice site variants with CADD c-scores >20

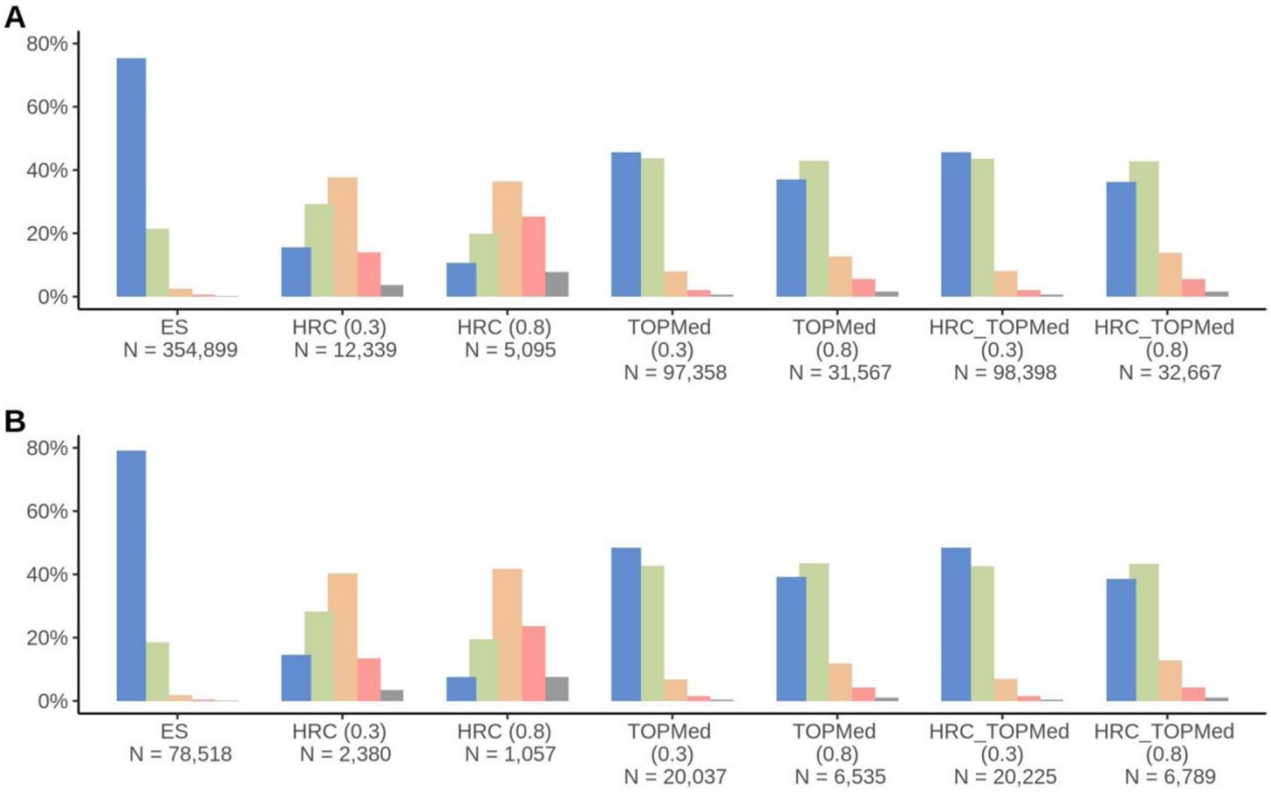


The minor allele frequency (MAF) categories are shown in the following colors: ***Blue*** (MAF$\leq 1\times10^{-5}$; ***Green*** ($1\times10^{-5}<$ MAF $\leq 1\times10^{-4}$); ***Orange*** ($1\times10^{-4}<$MAF$\leq 1\times10^{-3}$); ***Pink*** ($1\times10^{-3}<$ MAF$\leq 5\times10^{-3}$); and ***Grey*** ($5\times10^{-3}<$ MAF$<1\times10^{-2}$). The numbers in brackets represent the $R^2$ threshold and N is the number of variants in each MAF category. In the panels are displayed variants type: **(A)** Missense variants with CADD_c-score >= 20, **(B)** Splice site variants with CADD_c-score >= 20. For details on the percentage of variants in each MAF category, refer to Table S9. **Abbreviations**: CADD, combined annotation–dependent depletion; ES, exome sequence; HRC, Haplotype Reference Consortium; LoF, loss of function; and TOPMed, Trans-Omics for Precision Medicine.

**Supplementary Figure 2.** Comparison of unique variants for ES, TOPMed, and HRC datasets

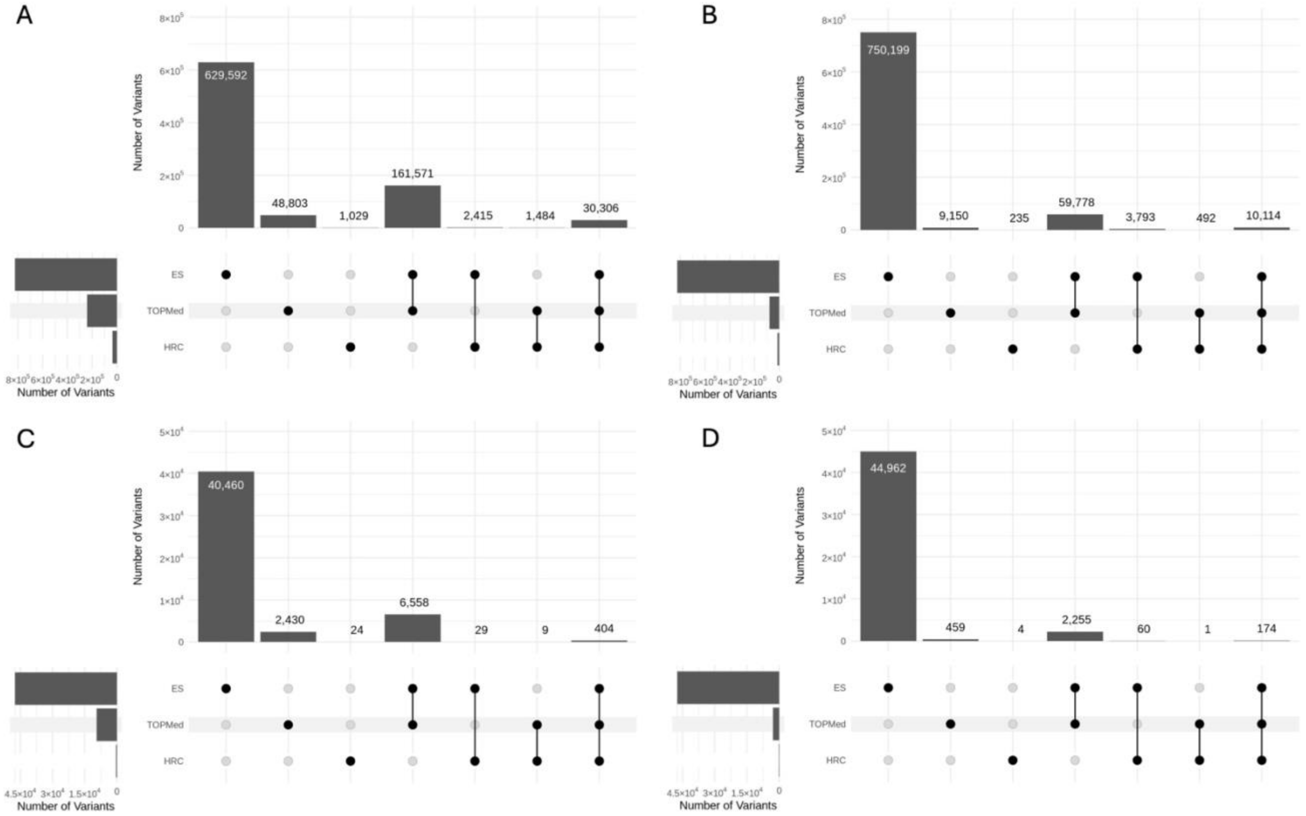


The upset plot displays the number of variants which are unique to each dataset as well as variants which are uniquely shared between two or more datasets. The variants included in the upset plot have a sample MAF of < 0.01 and include: **(A)** all LoF, missense, and splice site variants ($R^2 > 0.3$); **(B)** all variants ($R^2 > 0.8$); **(C)** LoF, missense, and splice site variants with CADD_c-score >= 20 ($R^2 > 0.3$); **(D)** LoF, missense, and splice site variants with CADD_c-score >= 20 ($R^2 > 0.8$). **Abbreviations**: ES: exome sequence; HRC: Haplotype Reference Consortium; LoF: loss of function; MAF: minor allele frequency; and TOPMed: Trans-Omics for Precision Medicine.

**Supplementary Figure 3.** Quantile-quantile plots of Type I error for exome sequence and imputed data

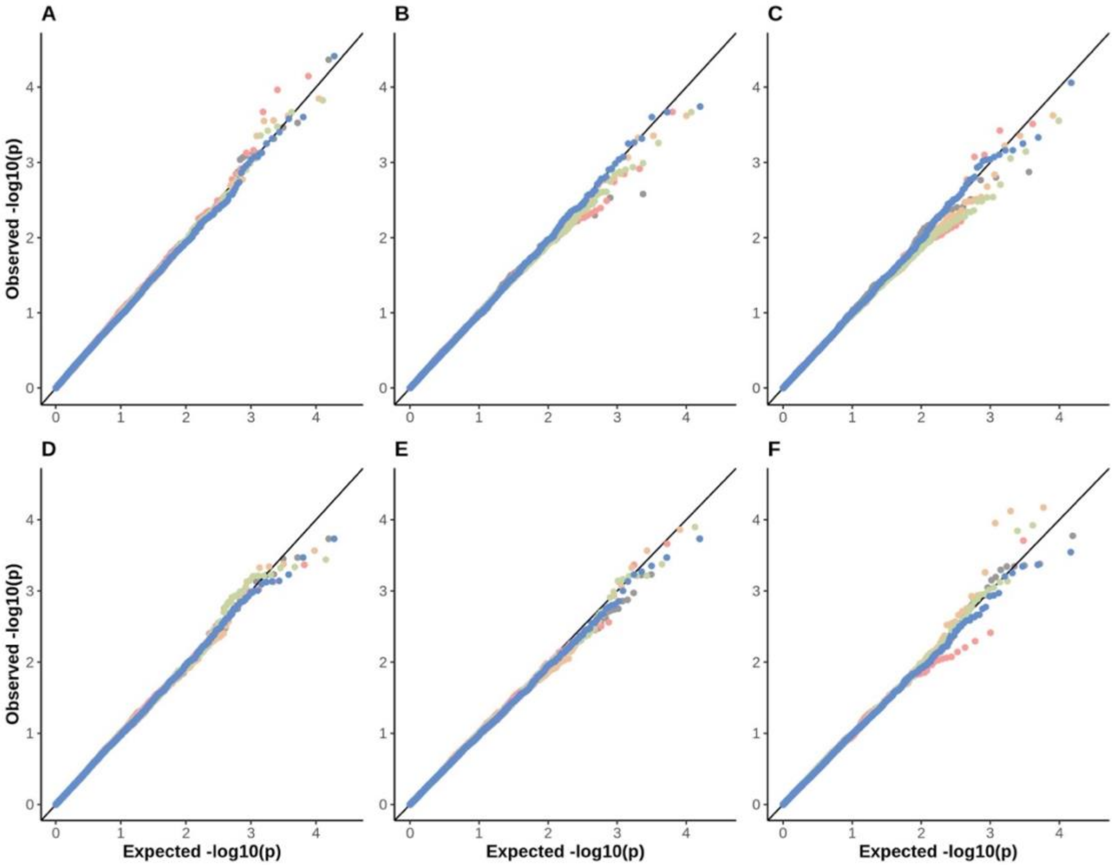


Type I error was evaluated by generating cases (N=40,000) and controls (N=60,000) under the null for each gene on chromosomes 1 and 2 with at least two variants. The data type is represented by the following colors: ***Pink*** (ES); ***Orange*** (HRC); ***Green*** (TOPMed); ***Blue*** (HRC_TOPMed); and ***Grey*** (ES_HRC_TOPMed). The $R^2$ value for imputed variant for panels: **A-C** $R^2>0.3$ and **D-F** $R^2>0.8$. The following rare variant MAF thresholds were used for the analysis MAF<0.01 panels **A** and **D**; MAF<0.005 panels **B** and **E**; and MAF<0.001 panels **C** and **F**. **Abbreviations**: ES: exome sequence; HRC: Haplotype Reference Consortium; MAF: minor allele frequency; and TOPMed: Trans-Omics for Precision Medicine.

**Supplementary Figure 4:** Power to detect rare-variant aggregate associations for exome sequence and imputed variant data

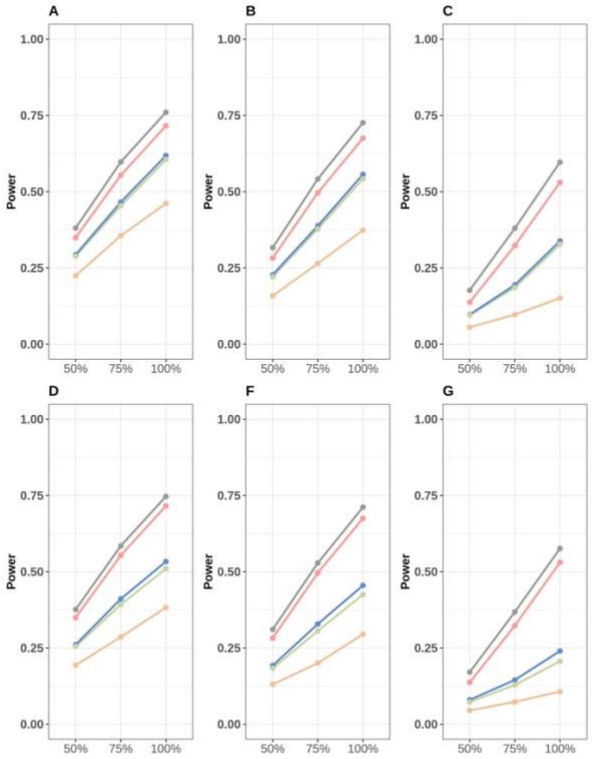


Power was evaluated by generating cases (N=40,000) and controls (N=60,000) for each gene on chromosomes 1 and 2 with at least two variants. The disease prevalence was 0.1, and the proportion of causal rare variants with each gene region varied between 100% and 50%, with each causal rare variant within a gene region having an odds ratio of 1.5. Power was estimated as the number of genes that met an exome-wide significance level of $2.5\times10^{-6}$ divided by the maximum number of genes tested for ES, HRC, TOPMED, HRC_TOPMed, or ES_HRC_TOPMed. The data type is represented by the following colors: ***Pink*** (ES); ***Orange*** (HRC); ***Green*** (TOPMed); ***Blue*** (HRC_TOPMed); and ***Grey*** (ES_HRC_TOPMed). The $R^2$ value for the imputed variant for panels **A-C** $R^2>0.3$ and **D-F** $R^2>0.8$. The following rare variant MAF thresholds were used for the analysis MAF<0.01 panels **A** and **D**; MAF<0,005 panels **B** and **E**; and MAF<0.001 panels **C** and **F**. **Abbreviations**: ES, exome sequence; HRC: Haplotype Reference Consortium; MAF, minor allele frequency; and TOPMed: Trans-Omics for Precision Medicine.

**Supplementary Table 1.** Number of variants for each rare variant minor allele frequency threshold and their mean minor allele frequencies

| | | | MAF < 0.01 | | MAF < 0.005 | | MAF < 0.001 | |
|---|---|---|---|---|---|---|---|---|
| **Dataset** | **$R^2$** | **Category** | **# Var (# Var)[a]** | **μ MAF(μ MAF)[b]** | **# Var (# Var)[a]** | **μ MAF(μ MAF)[b]** | **# Var (# Var)[a]** | **μ MAF(μ MAF)[b]** |
| HRC_TOPMed | >0.3 | LoF | 11,030 | $1.25\times10^{-4}$ | 10,992 | $9.23\times10^{-5}$ | 10,836 | $5.16\times10^{-5}$ |
| | | Missense | 169,812 (98,398) | $1.70\times10^{-4}$ ($1.49\times10^{-4}$) | 168,813 (97,921) | $1.23\times10^{-4}$ ($1.10\times10^{-4}$) | 164,753 (95,868) | $6.14\times10^{-5}$ ($5.71\times10^{-5}$) |
| | | Splice site | 64,820 (20,225) | $1.85\times10^{-4}$ ($1.34\times10^{-4}$) | 64,420 (20,147) | $1.29\times10^{-4}$ ($1.00\times10^{-4}$) | 62,911 (19,791) | $6.26\times10^{-5}$ ($5.38\times10^{-5}$) |
| | | All | 245,663 | $1.72\times10^{-4}$ | 244,225 | $1.23\times10^{-4}$ | 238,500 | $6.12\times10^{-5}$ |
| | >0.8 | LoF | 3,485 | $2.65\times10^{-4}$ | 3,451 | $1.84\times10^{-4}$ | 3,319 | $7.96\times10^{-5}$ |
| | | Missense | 57,643 (32,667) | $3.83\times10^{-4}$ ($3.39\times10^{-4}$) | 56,681 (32,206) | $2.56\times10^{-4}$ ($2.32\times10^{-4}$) | 52,981 (30,308) | $9.52\times10^{-5}$ ($8.90\times10^{-5}$) |
| | | Splice site | 22,611 (6,789) | $3.88\times10^{-4}$ ($2.99\times10^{-4}$) | 22,251 (6,712) | $2.52\times10^{-4}$ ($2.06\times10^{-4}$) | 20,982 (6,389) | $9.42\times10^{-5}$ ($8.52\times10^{-5}$) |
| | | All | 83,739 | $3.80\times10^{-4}$ | 82,383 | $2.52\times10^{-4}$ | 77,282 | $9.43\times10^{-5}$ |
| ES_HRC_TOPMed | >0.3 | LoF | 56,193 | $6.67\times10^{-5}$ | 56,147 | $5.09\times10^{-5}$ | 55,953 | $3.04\times10^{-5}$ |
| | | Missense | 600,709 (355,228) | $9.06\times10^{-5}$ ($7.93\times10^{-5}$) | 599,635 (354,738) | $6.64\times10^{-5}$ ($6.00\times10^{-5}$) | 595,348 (352,631) | $3.60\times10^{-5}$ ($3.39\times10^{-5}$) |
| | | Splice site | 218,327 (78,612) | $1.02\times\times10^{-4}$ ($7.10\times10^{-5}$) | 217,882 (78,535) | $7.23\times10^{-5}$ ($5.53\times10^{-5}$) | 216,243 (78,156) | $3.73\times10^{-5}$ ($3.20\times10^{-5}$) |
| | | All | 875,229 | $9.22\times10^{-5}$ | 873,664 | $6.71\times10^{-5}$ | 867,544 | $3.60\times10^{-5}$ |
| | >0.8 | LoF | 53,822 | $6.88\times10^{-5}$ | 53,778 | $5.30\times10^{-5}$ | 53,586 | $3.14\times10^{-5}$ |
| | | Missense | 575,036 (354,919) | $9.40\times10^{-5}$ ($7.85\times10^{-5}$) | 573,972 (354,438) | $6.88\times10^{-5}$ ($5.96\times10^{-5}$) | 569,733 (352,365) | $3.72\times10^{-5}$ ($3.39\times10^{-5}$) |
| | | Splice site | 204,931 (78,536) | $1.06\times10^{-4}$ ($7.06\times10^{-5}$) | 204,494 (78,460) | $7.50\times10^{-5}$ ($5.51\times10^{-5}$) | 202,885 (78,085) | $3.85\times10^{-5}$ ($3.20\times10^{-5}$) |
| | | All | 833,789 | $9.57\times10^{-5}$ | 832,244 | $6.95\times10^{-5}$ | 826,204 | $3.72\times10^{-5}$ |

The MAF for each variant is obtained from gnomAD for NFE. [a] Number of variants (Number of variants with CADD c-score ≥ 20). [b] Mean MAF (Mean MAF of variants with CADD c-score ≥ 20). **Abbreviations:** # Var, number of variants; μ, mean; CADD, combined annotation–dependent depletion; ES, exome sequence; gnomAD, Genome Aggregation Database; HRC, Haplotype Reference Consortium; LoF, loss of function; MAF, minor allele frequency; NFE, non-Finnish Europeans; $R^2$, imputation quality; and TOPMed, Trans-Omics for Precision Medicine.

**Supplementary Table 2.** Welch two sample T-test comparing mean $R^2$ values for imputed variant datasets

| Dataset | MAF | p-value [a] |
|---|---|---|
| HRC v.s. TOPMed | MAF < 0.001 | $6.46\times10^{-67}$ |
| | MAF < 0.001[b] | $1.87\times10^{-63}$ |
| | 0.001 ≤ MAF < 0.005 | $8.48\times10^{-17}$ |
| | 0.001 ≤ MAF < 0.005[b] | $2.07\times10^{-18}$ |
| | 0.005 ≤ MAF < 0.01 | $2.48\times10^{-7}$ |
| | 0.005 ≤ MAF < 0.01[b] | $1.68\times10^{-9}$ |
| HRC v.s. HRC_TOPMed | MAF < 0.001 | $1.18\times10^{-36}$ |
| | 0.001 ≤ MAF < 0.005 | $3.62\times10^{-52}$ |
| | 0.005 ≤ MAF < 0.01 | $4.34\times10^{-5}$ |
| TOPMed v.s. HRC_TOPMed | MAF < 0.001 | $<2.23\times10^{-308}$ |
| | 0.001 ≤ MAF < 0.005 | $3.16\times10^{-3}$ |
| | 0.005 ≤ MAF < 0.01 | $4.34\times10^{-5}$ |

All LoF, missense, and splice site variants obtained from chromosomes 1 and 2 were included. The MAF for each variant is obtained from gnomAD for NFE. [a]A two-sided test was performed. [b]Includes only the imputed variants that overlap between HRC and TOPMed. **Abbreviations:** gnomAD, Genome Aggregation Database; HRC, Haplotype Reference Consortium; LoF, loss of function; MAF, minor allele frequency; NFE, non-Finnish Europeans; $R^2$, imputation quality; and TOPMed, Trans-Omics for Precision Medicine.

**Supplementary Table 3.** Welch Two sample T-test comparing mean $R^2$ and mean $r^2$ for imputed variants

| **Dataset** | **MAF** | **p-value [a]** |
|---|---|---|
| HRC | MAF < 0.001 | $1.96\times10^{-20}$ |
| | MAF < 0.001[b] | $8.89\times10^{-15}$ |
| | 0.001 ≤ MAF < 0.005 | $4.46\times10^{-16}$ |
| | 0.001 ≤ MAF < 0.005[b] | $4.50\times10^{-19}$ |
| | 0.005 ≤ MAF < 0.01 | $2.29\times10^{-1}$ |
| | 0.005 ≤ MAF < 0.01[b] | $5.69\times10^{-1}$ |
| TOPMed | MAF < 0.001 | $9.95\times10^{-63}$ |
| | MAF<0.001[b] | $<2.23\times10^{-308}$ |
| | 0.001 ≤ MAF < 0.005 | $1.24\times10^{-53}$ |
| | 0.001 ≤ MAF < 0.005[b] | $8.56\times10^{-85}$ |
| | 0.005 ≤ MAF < 0.01 | $8.00\times10^{-1}$ |
| | 0.005 ≤ MAF < 0.01[b] | $8.28\times10^{-1}$ |
| HRC_TOPMed | MAF < 0.001 | $6.12\times10^{-1}$ |
| | 0.001 ≤ MAF < 0.005 | $4.59\times10^{-12}$ |
| | 0.005 ≤ MAF < 0.01 | $1.15\times10^{-188}$ |

All LoF, missense, and splice site variants obtained from chromosomes 1 and 2 were included. The MAF for each variant is obtained from gnomAD for NFE. [a] A two-sided test was performed. [b] Includes only the imputed variants that overlap between HRC and TOPMed. **Abbreviations:** gnomAD, Genome Aggregation Database; HRC, Haplotype Reference Consortium; LoF, loss of function; MAF, minor allele frequency; NFE, non-Finnish Europeans; $r^2$, correlation between imputed and exome sequence data variants; $R^2$, imputation quality; and TOPMed, Trans-Omics for Precision Medicine.

**Supplementary Table 4.** Power to detect rare-variant aggregate associations for exome sequence and imputed variant ($R^2 > 0.3$)

| Prevalence | Effect size | Deleterious Proportion | Power | | | | |
|---|---|---|---|---|---|---|---|
| | | | ES | ES_HRC_TOPMed | HRC_TOPMed | HRC | TOPMed |
| **MAF < 0.01** | | | | | | | |
| 0.1 | 1.2 | 0.5 | 0.054 | 0.085 | 0.065 | 0.050 | 0.063 |
| | | 0.75 | 0.138 | 0.179 | 0.136 | 0.095 | 0.130 |
| | | 1 | 0.250 | 0.313 | 0.226 | 0.166 | 0.220 |
| | 1.5 | 0.5 | 0.350 | 0.381 | 0.293 | 0.225 | 0.289 |
| | | 0.75 | 0.554 | 0.598 | 0.466 | 0.356 | 0.454 |
| | | 1 | 0.716 | 0.761 | 0.618 | 0.461 | 0.605 |
| | 1.8 | 0.5 | 0.505 | 0.528 | 0.415 | 0.312 | 0.404 |
| | | 0.75 | 0.703 | 0.741 | 0.586 | 0.441 | 0.576 |
| | | 1 | 0.810 | 0.848 | 0.707 | 0.520 | 0.694 |
| 0.2 | 1.2 | 0.5 | 0.058 | 0.091 | 0.067 | 0.052 | 0.062 |
| | | 0.75 | 0.148 | 0.196 | 0.145 | 0.104 | 0.140 |
| | | 1 | 0.266 | 0.326 | 0.242 | 0.175 | 0.230 |
| | 1.5 | 0.5 | 0.351 | 0.383 | 0.292 | 0.222 | 0.284 |
| | | 0.75 | 0.562 | 0.608 | 0.470 | 0.354 | 0.456 |
| | | 1 | 0.721 | 0.771 | 0.627 | 0.470 | 0.615 |
| | 1.8 | 0.5 | 0.505 | 0.530 | 0.418 | 0.311 | 0.411 |
| | | 0.75 | 0.707 | 0.739 | 0.594 | 0.437 | 0.582 |
| | | 1 | 0.805 | 0.845 | 0.704 | 0.516 | 0.688 |
| **MAF < 0.005** | | | | | | | |
| 0.1 | 1.2 | 0.5 | 0.036 | 0.069 | 0.053 | 0.040 | 0.051 |
| | | 0.75 | 0.086 | 0.131 | 0.096 | 0.064 | 0.090 |
| | | 1 | 0.177 | 0.243 | 0.159 | 0.108 | 0.155 |
| | 1.5 | 0.5 | 0.283 | 0.317 | 0.228 | 0.159 | 0.221 |
| | | 0.75 | 0.496 | 0.542 | 0.388 | 0.265 | 0.378 |
| | | 1 | 0.675 | 0.726 | 0.557 | 0.374 | 0.542 |
| | 1.8 | 0.5 | 0.470 | 0.498 | 0.365 | 0.243 | 0.355 |
| | | 0.75 | 0.677 | 0.720 | 0.541 | 0.378 | 0.530 |
| | | 1 | 0.787 | 0.831 | 0.665 | 0.452 | 0.656 |
| 0.2 | 1.2 | 0.5 | 0.035 | 0.066 | 0.050 | 0.038 | 0.046 |
| | | 0.75 | 0.097 | 0.147 | 0.104 | 0.073 | 0.101 |

| Prevalence | Effect size | Deleterious Proportion | Power | | | | |
|---|---|---|---|---|---|---|---|
| | | | ES | ES_HRC_TOPMed | HRC_TOPMed | HRC | TOPMed |
| 0.2 | 1.2 | 1 | 0.201 | 0.261 | 0.174 | 0.112 | 0.164 |
| | 1.5 | 0.5 | 0.287 | 0.321 | 0.220 | 0.156 | 0.216 |
| | | 0.75 | 0.510 | 0.556 | 0.400 | 0.275 | 0.387 |
| | | 1 | 0.678 | 0.734 | 0.563 | 0.380 | 0.553 |
| | 1.8 | 0.5 | 0.468 | 0.496 | 0.365 | 0.249 | 0.360 |
| | | 0.75 | 0.680 | 0.717 | 0.548 | 0.364 | 0.537 |
| | | 1 | 0.780 | 0.825 | 0.663 | 0.446 | 0.650 |
| **MAF < 0.001** | | | | | | | |
| 0.1 | 1.2 | 0.5 | 0.022 | 0.055 | 0.044 | 0.033 | 0.043 |
| | | 0.75 | 0.042 | 0.089 | 0.068 | 0.046 | 0.065 |
| | | 1 | 0.090 | 0.154 | 0.096 | 0.064 | 0.092 |
| | 1.5 | 0.5 | 0.137 | 0.177 | 0.097 | 0.055 | 0.096 |
| | | 0.75 | 0.323 | 0.380 | 0.195 | 0.097 | 0.186 |
| | | 1 | 0.531 | 0.597 | 0.338 | 0.151 | 0.328 |
| | 1.8 | 0.5 | 0.331 | 0.374 | 0.189 | 0.086 | 0.185 |
| | | 0.75 | 0.585 | 0.637 | 0.364 | 0.169 | 0.359 |
| | | 1 | 0.710 | 0.762 | 0.510 | 0.237 | 0.504 |
| 0.2 | 1.2 | 0.5 | 0.022 | 0.054 | 0.044 | 0.031 | 0.041 |
| | | 0.75 | 0.050 | 0.098 | 0.071 | 0.050 | 0.068 |
| | | 1 | 0.099 | 0.162 | 0.103 | 0.066 | 0.098 |
| | 1.5 | 0.5 | 0.139 | 0.181 | 0.099 | 0.056 | 0.096 |
| | | 0.75 | 0.335 | 0.397 | 0.204 | 0.100 | 0.194 |
| | | 1 | 0.541 | 0.611 | 0.344 | 0.150 | 0.337 |
| | 1.8 | 0.5 | 0.322 | 0.362 | 0.186 | 0.086 | 0.183 |
| | | 0.75 | 0.580 | 0.632 | 0.376 | 0.158 | 0.367 |
| | | 1 | 0.711 | 0.765 | 0.516 | 0.228 | 0.504 |

Power was estimated by generating cases (N=40,000) and controls (N=60,000) for each gene on chromosomes 1 and 2 with at least two variants and is calculated as the number of genes that met an exome-wide significance level of $2.5\times10^{-6}$ divided by the maximum number of genes tested for ES, HRC, TOPMED, HRC_TOPMed, or ES_HRC_TOPMed. **Abbreviations:** ES, exome sequence; HRC, Haplotype Reference Consortium; MAF, minor allele frequency; $R^2$, imputation quality; and TOPMed, Trans-Omics for Precision Medicine.

**Supplementary Table 5.** Power to detect rare-variant aggregate associations for exome sequence and imputed variant ($R^2 > 0.8$)

| Prevalence | Effect size | Deleterious Proportion | Power | | | | |
|---|---|---|---|---|---|---|---|
| | | | ES | ES_HRC_TOPMed | HRC_TOPMed | HRC | TOPMed |
| **MAF < 0.01** | | | | | | | |
| 0.1 | 1.2 | 0.5 | 0.054 | 0.082 | 0.058 | 0.047 | 0.059 |
| | | 0.75 | 0.138 | 0.175 | 0.122 | 0.087 | 0.119 |
| | | 1 | 0.250 | 0.304 | 0.203 | 0.142 | 0.191 |
| | 1.5 | 0.5 | 0.350 | 0.377 | 0.261 | 0.194 | 0.255 |
| | | 0.75 | 0.554 | 0.585 | 0.411 | 0.286 | 0.392 |
| | | 1 | 0.716 | 0.747 | 0.534 | 0.383 | 0.510 |
| | 1.8 | 0.5 | 0.505 | 0.523 | 0.366 | 0.259 | 0.346 |
| | | 0.75 | 0.703 | 0.727 | 0.494 | 0.351 | 0.468 |
| | | 1 | 0.810 | 0.836 | 0.594 | 0.421 | 0.567 |
| 0.2 | 1.2 | 0.5 | 0.058 | 0.087 | 0.062 | 0.048 | 0.058 |
| | | 0.75 | 0.148 | 0.192 | 0.134 | 0.092 | 0.126 |
| | | 1 | 0.266 | 0.320 | 0.216 | 0.155 | 0.203 |
| | 1.5 | 0.5 | 0.351 | 0.376 | 0.259 | 0.189 | 0.250 |
| | | 0.75 | 0.562 | 0.594 | 0.415 | 0.294 | 0.394 |
| | | 1 | 0.721 | 0.756 | 0.539 | 0.385 | 0.515 |
| | 1.8 | 0.5 | 0.505 | 0.521 | 0.354 | 0.256 | 0.343 |
| | | 0.75 | 0.707 | 0.729 | 0.501 | 0.351 | 0.477 |
| | | 1 | 0.805 | 0.831 | 0.590 | 0.413 | 0.562 |
| **MAF < 0.005** | | | | | | | |
| 0.1 | 1.2 | 0.5 | 0.036 | 0.066 | 0.047 | 0.037 | 0.046 |
| | | 0.75 | 0.086 | 0.127 | 0.084 | 0.058 | 0.081 |
| | | 1 | 0.177 | 0.236 | 0.139 | 0.091 | 0.130 |
| | 1.5 | 0.5 | 0.283 | 0.311 | 0.192 | 0.131 | 0.183 |
| | | 0.75 | 0.496 | 0.529 | 0.329 | 0.200 | 0.306 |
| | | 1 | 0.675 | 0.712 | 0.455 | 0.295 | 0.425 |
| | 1.8 | 0.5 | 0.470 | 0.491 | 0.304 | 0.187 | 0.283 |
| | | 0.75 | 0.677 | 0.704 | 0.436 | 0.280 | 0.406 |
| | | 1 | 0.787 | 0.817 | 0.537 | 0.342 | 0.501 |
| 0.2 | 1.2 | 0.5 | 0.035 | 0.064 | 0.047 | 0.035 | 0.044 |
| | | 0.75 | 0.097 | 0.144 | 0.095 | 0.063 | 0.089 |

| Prevalence | Effect size | Deleterious Proportion | Power | | | | |
|---|---|---|---|---|---|---|---|
| | | | ES | ES_HRC_TOPMed | HRC_TOPMed | HRC | TOPMed |
| | | 1 | 0.201 | 0.257 | 0.150 | 0.097 | 0.140 |
| | 1.5 | 0.5 | 0.287 | 0.315 | 0.187 | 0.130 | 0.176 |
| | | 0.75 | 0.510 | 0.543 | 0.336 | 0.213 | 0.311 |
| | | 1 | 0.678 | 0.718 | 0.462 | 0.296 | 0.431 |
| | 1.8 | 0.5 | 0.468 | 0.488 | 0.299 | 0.191 | 0.282 |
| | | 0.75 | 0.680 | 0.705 | 0.442 | 0.278 | 0.413 |
| | | 1 | 0.780 | 0.810 | 0.532 | 0.333 | 0.496 |
| **MAF < 0.001** | | | | | | | |
| 0.1 | 1.2 | 0.5 | 0.022 | 0.053 | 0.041 | 0.030 | 0.040 |
| | | 0.75 | 0.042 | 0.086 | 0.059 | 0.041 | 0.059 |
| | | 1 | 0.090 | 0.149 | 0.089 | 0.057 | 0.085 |
| | 1.5 | 0.5 | 0.137 | 0.172 | 0.080 | 0.046 | 0.074 |
| | | 0.75 | 0.323 | 0.371 | 0.146 | 0.074 | 0.130 |
| | | 1 | 0.531 | 0.580 | 0.242 | 0.108 | 0.208 |
| | 1.8 | 0.5 | 0.331 | 0.360 | 0.139 | 0.066 | 0.122 |
| | | 0.75 | 0.585 | 0.625 | 0.250 | 0.117 | 0.215 |
| | | 1 | 0.710 | 0.749 | 0.342 | 0.150 | 0.295 |
| 0.2 | 1.2 | 0.5 | 0.022 | 0.052 | 0.040 | 0.028 | 0.039 |
| | | 0.75 | 0.050 | 0.096 | 0.066 | 0.045 | 0.063 |
| | | 1 | 0.099 | 0.161 | 0.091 | 0.057 | 0.087 |
| | 1.5 | 0.5 | 0.139 | 0.174 | 0.082 | 0.047 | 0.076 |
| | | 0.75 | 0.335 | 0.381 | 0.146 | 0.075 | 0.130 |
| | | 1 | 0.541 | 0.595 | 0.234 | 0.107 | 0.206 |
| | 1.8 | 0.5 | 0.322 | 0.348 | 0.133 | 0.061 | 0.118 |
| | | 0.75 | 0.580 | 0.622 | 0.251 | 0.108 | 0.215 |
| | | 1 | 0.711 | 0.750 | 0.340 | 0.153 | 0.292 |

Power was estimated by generating cases (N=40,000) and controls (N=60,000) for each gene on chromosomes 1 and 2 with at least two variants and is calculated as the number of genes that met an exome-wide significance level of $2.5\times10^{-6}$ divided by the maximum number of genes tested for ES, HRC, TOPMED, HRC_TOPMed, or ES_HRC_TOPMed. **Abbreviations:** ES, exome sequence; HRC, Haplotype Reference Consortium; MAF, minor allele frequency; $R^2$, imputation quality; and TOPMed, Trans-Omics for Precision Medicine.

**Supplementary Table 6.** Percentage change in power compared to exome sequence data for imputed datasets ($R^2 > 0.3$)

| Prevalence | Effect size | Percentage change in power compared to ES | | |
|---|---|---|---|---|
| | | HRC_TOPMed | HRC | TOPMed |
| **MAF < 0.01** | | | | |
| 0.1 | 1.2 | -9.49% | -33.46% | -11.99% |
| | 1.5 | -13.61% | -35.54% | -15.44% |
| | 1.8 | -12.71% | -35.90% | -14.41% |
| 0.2 | 1.2 | -9.04% | -34.04% | -13.50% |
| | 1.5 | -13.08% | -34.78% | -14.68% |
| | 1.8 | -12.49% | -35.92% | -14.55% |
| **MAF < 0.05** | | | | |
| 0.1 | 1.2 | -10.23% | -38.80% | -12.70% |
| | 1.5 | -17.57% | -44.66% | -19.74% |
| | 1.8 | -15.58% | -42.62% | -16.65% |
| 0.2 | 1.2 | -13.66% | -44.41% | -18.48% |
| | 1.5 | -16.95% | -43.90% | -18.47% |
| | 1.8 | -15.01% | -42.83% | -16.61% |
| **MAF < 0.001** | | | | |
| 0.1 | 1.2 | 6.92% | -29.41% | 2.08% |
| | 1.5 | -36.24% | -71.53% | -38.29% |
| | 1.8 | -28.07% | -66.65% | -29.04% |
| 0.2 | 1.2 | 3.46% | -33.65% | -0.94% |
| | 1.5 | -36.47% | -72.30% | -37.62% |
| | 1.8 | -27.36% | -67.90% | -29.12% |

Variants were obtained from chromosomes 1 and 2. Percentage change in power was calculated based on simulation results where all variants were causal. The proportion of causal rare variants with each gene region was 100%. **Abbreviations:** CADD, combined annotation–dependent depletion; ES, exome sequence; HRC, Haplotype Reference Consortium; LoF, loss of function; MAF, minor allele frequency; $R^2$, imputation quality; and TOPMed, Trans-Omics for Precision Medicine.

**Supplementary Table 7.** Percentage change in power compared to exome sequence data for imputed datasets ($R^2 > 0.8$)

| Prevalence | Effect size | Percentage change in power compared to ES | | |
|---|---|---|---|---|
| | | **HRC_TOPMed** | **HRC** | **TOPMed** |
| | | **MAF < 0.01** | | |
| 0.1 | 1.2 | -18.73% | -43.07% | -23.47% |
| | 1.5 | -25.47% | -46.53% | -28.70% |
| | 1.8 | -26.69% | -48.07% | -30.08% |
| 0.2 | 1.2 | -18.66% | -41.90% | -23.59% |
| | 1.5 | -25.25% | -46.60% | -28.63% |
| | 1.8 | -26.69% | -48.64% | -30.18% |
| | | **MAF < 0.05** | | |
| 0.1 | 1.2 | -21.52% | -48.68% | -26.63% |
| | 1.5 | -32.59% | -56.26% | -37.03% |
| | 1.8 | -31.84% | -56.62% | -36.36% |
| 0.2 | 1.2 | -25.47% | -51.71% | -30.28% |
| | 1.5 | -31.83% | -56.33% | -36.48% |
| | 1.8 | -31.75% | -57.33% | -36.43% |
| | | **MAF < 0.001** | | |
| 0.1 | 1.2 | -1.75% | -36.94% | -5.58% |
| | 1.5 | -54.45% | -79.68% | -60.79% |
| | 1.8 | -51.80% | -78.83% | -58.45% |
| 0.2 | 1.2 | -8.17% | -42.37% | -12.29% |
| | 1.5 | -56.71% | -80.30% | -61.88% |
| | 1.8 | -52.15% | -78.42% | -58.92% |

Variants were obtained from chromosomes 1 and 2. Percentage change in power was calculated based on simulation results where all variants were causal. The proportion of causal rare variants with each gene region was 100%. **Abbreviations:** CADD, combined annotation–dependent depletion; ES, exome sequence; HRC, Haplotype Reference Consortium; LoF, loss of function; MAF, minor allele frequency; $R^2$, imputation quality; and TOPMed, Trans-Omics for Precision Medicine.

**Supplementary Table 8A.** Variant Minor Allele Frequency Plot Statistics for Figure 1A

| Data | $R^2$ (>) | MAF category | Proportion |
|---|---|---|---|
| ES | - | $0.005 < MAF < 0.01$ | 0.20% |
| | | $0.001 < MAF \leq 0.005$ | 0.70% |
| | | $1\times10^{-4} < MAF \leq 0.001$ | 2.50% |
| | | $1\times10^{-5} < MAF \leq 1\times10^{-4}$ | 21.40% |
| | | $MAF \leq 1\times10^{-5}$ | 75.30% |
| HRC | 0.3 | $0.005 < MAF < 0.01$ | 3.70% |
| | | $0.001 < MAF \leq 0.005$ | 14% |
| | | $1\times10^{-4} < MAF \leq 0.001$ | 37.70% |
| | | $1\times10^{-5} < MAF \leq 1\times10^{-4}$ | 29.20% |
| | | $MAF \leq 1\times10^{-5}$ | 15.50% |
| | 0.8 | $0.005 < MAF < 0.01$ | 7.80% |
| | | $0.001 < MAF \leq 0.005$ | 25.30% |
| | | $1\times10^{-4} < MAF \leq 0.001$ | 36.40% |
| | | $1\times10^{-5} < MAF \leq 1\times10^{-4}$ | 19.80% |
| | | $MAF \leq 1\times10^{-5}$ | 10.70% |
| TOPMed | 0.3 | $0.005 < MAF < 0.01$ | 0.60% |
| | | $0.001 < MAF \leq 0.005$ | 2.10% |
| | | $1\times10^{-4} < MAF \leq 0.001$ | 8% |
| | | $1\times10^{-5} < MAF \leq 1\times10^{-4}$ | 43.70% |
| | | $MAF \leq 1\times10^{-5}$ | 45.70% |
| | 0.8 | $0.005 < MAF < 0.01$ | 1.60% |
| | | $0.001 < MAF \leq 0.005$ | 5.60% |
| | | $1\times10^{-4} < MAF \leq 0.001$ | 12.70% |
| | | $1\times10^{-5} < MAF \leq 1\times10^{-4}$ | 43% |
| | | $MAF \leq 1\times10^{-5}$ | 37.10% |
| HRC_TOPMed | 0.3 | $0.005 < MAF < 0.01$ | 0.60% |
| | | $0.001 < MAF \leq 0.005$ | 2.10% |
| | | $1\times10^{-4} < MAF \leq 0.001$ | 8.10% |
| | | $1\times10^{-5} < MAF \leq 1\times10^{-4}$ | 43.60% |
| | | $MAF \leq 1\times10^{-5}$ | 45.60% |
| | 0.8 | $0.005 < MAF < 0.01$ | 1.60% |
| | | $0.001 < MAF \leq 0.005$ | 5.60% |
| | | $1\times10^{-4} < MAF \leq 0.001$ | 13.80% |
| | | $1\times10^{-5} < MAF \leq 1\times10^{-4}$ | 42.80% |
| | | $MAF \leq 1\times10^{-5}$ | 36.20% |

Variants were obtained from chromosomes 1 and 2. The MAF for each variant is obtained from gnomAD for NFE. **Abbreviations**: ES, exome sequencing; gnomAD, Genome Aggregation Database; HRC, Haplotype Reference Consortium; MAF, minor allele frequency; NFE, non-Finnish European; $R^2$, imputation quality; and TOPMed, Trans-Omics for Precision Medicine.

**Supplementary Table 8B.** Variant Minor Allele Frequency Plot Statistics for Figure 1B

| Data | $R^2$ (>) | MAF category | Proportion |
|---|---|---|---|
| ES | - | $0.005 < MAF < 0.01$ | 0.10% |
| | | $0.001 < MAF \leq 0.005$ | 0.30% |
| | | $1\times10^{-4} < MAF \leq 0.001$ | 1.40% |
| | | $1\times10^{-5} < MAF \leq 1\times10^{-4}$ | 14.80% |
| | | $MAF \leq 1\times10^{-5}$ | 83.40% |
| HRC | 0.3 | $0.005 < MAF < 0.01$ | 3.50% |
| | | $0.001 < MAF \leq 0.005$ | 13.60% |
| | | $1\times10^{-4} < MAF \leq 0.001$ | 39% |
| | | $1\times10^{-5} < MAF \leq 1\times10^{-4}$ | 29.90% |
| | | $MAF \leq 1\times10^{-5}$ | 14% |
| | 0.8 | $0.005 < MAF < 0.01$ | 5.90% |
| | | $0.001 < MAF \leq 0.005$ | 20.80% |
| | | $1\times10^{-4} < MAF \leq 0.001$ | 44.10% |
| | | $1\times10^{-5} < MAF \leq 1\times10^{-4}$ | 23.30% |
| | | $MAF \leq 1\times10^{-5}$ | 5.90% |
| TOPMed | 0.3 | $0.005 < MAF < 0.01$ | 0.30% |
| | | $0.001 < MAF \leq 0.005$ | 1.30% |
| | | $1\times10^{-4} < MAF \leq 0.001$ | 5.80% |
| | | $1\times10^{-5} < MAF \leq 1\times10^{-4}$ | 40.50% |
| | | $MAF \leq 1\times10^{-5}$ | 52% |
| | 0.8 | $0.005 < MAF < 0.01$ | 0.90% |
| | | $0.001 < MAF \leq 0.005$ | 3.60% |
| | | $1\times10^{-4} < MAF \leq 0.001$ | 9.80% |
| | | $1\times10^{-5} < MAF \leq 1\times10^{-4}$ | 42.80% |
| | | $MAF \leq 1\times10^{-5}$ | 42.90% |
| HRC_TOPMed | 0.3 | $0.005 < MAF < 0.01$ | 0.30% |
| | | $0.001 < MAF \leq 0.005$ | 1.40% |
| | | $1\times10^{-4} < MAF \leq 0.001$ | 5.90% |
| | | $1\times10^{-5} < MAF \leq 1\times10^{-4}$ | 40.50% |
| | | $MAF \leq 1\times10^{-5}$ | 52% |
| | 0.8 | $0.005 < MAF < 0.01$ | 0.90% |
| | | $0.001 < MAF \leq 0.005$ | 3.70% |
| | | $1\times10^{-4} < MAF \leq 0.001$ | 10.80% |
| | | $1\times10^{-5} < MAF \leq 1\times10^{-4}$ | 42.50% |
| | | $MAF \leq 1\times10^{-5}$ | 42.20% |

Variants were obtained from chromosomes 1 and 2. The MAF for each variant is obtained from gnomAD for NFE. **Abbreviations**: ES, exome sequencing; gnomAD, Genome Aggregation Database; HRC, Haplotype Reference Consortium; MAF, minor allele frequency; NFE, non-Finnish European; $R^2$, imputation quality; and TOPMed, Trans-Omics for Precision Medicine.

**Supplementary Table 8C.** Variant Minor Allele Frequency Plot Statistics for Figure 1C

| Data | $R^2$ (>) | MAF category | Proportion |
|---|---|---|---|
| ES | - | $0.005 < MAF < 0.01$ | 0.20% |
| | | $0.001 < MAF \leq 0.005$ | 0.70% |
| | | $1\times10^{-4} < MAF \leq 0.001$ | 2.60% |
| | | $1\times10^{-5} < MAF \leq 1\times10^{-4}$ | 22.30% |
| | | $MAF \leq 1\times10^{-5}$ | 74.30% |
| HRC | 0.3 | $0.005 < MAF < 0.01$ | 3.80% |
| | | $0.001 < MAF \leq 0.005$ | 14.60% |
| | | $1\times10^{-4} < MAF \leq 0.001$ | 39.10% |
| | | $1\times10^{-5} < MAF \leq 1\times10^{-4}$ | 29.80% |
| | | $MAF \leq 1\times10^{-5}$ | 12.70% |
| | 0.8 | $0.005 < MAF < 0.01$ | 8% |
| | | $0.001 < MAF \leq 0.005$ | 27.40% |
| | | $1\times10^{-4} < MAF \leq 0.001$ | 37.80% |
| | | $1\times10^{-5} < MAF \leq 1\times10^{-4}$ | 19.40% |
| | | $MAF \leq 1\times10^{-5}$ | 7.40% |
| TOPMed | 0.3 | $0.005 < MAF < 0.01$ | 0.60% |
| | | $0.001 < MAF \leq 0.005$ | 2.10% |
| | | $1\times10^{-4} < MAF \leq 0.001$ | 8.20% |
| | | $1\times10^{-5} < MAF \leq 1\times10^{-4}$ | 45.40% |
| | | $MAF \leq 1\times10^{-5}$ | 43.70% |
| | 0.8 | $0.005 < MAF < 0.01$ | 1.60% |
| | | $0.001 < MAF \leq 0.005$ | 5.90% |
| | | $1\times10^{-4} < MAF \leq 0.001$ | 13.20% |
| | | $1\times10^{-5} < MAF \leq 1\times10^{-4}$ | 44.60% |
| | | $MAF \leq 1\times10^{-5}$ | 34.70% |
| HRC_TOPMed | 0.3 | $0.005 < MAF < 0.01$ | 0.60% |
| | | $0.001 < MAF \leq 0.005$ | 2.20% |
| | | $1\times10^{-4} < MAF \leq 0.001$ | 8.30% |
| | | $1\times10^{-5} < MAF \leq 1\times10^{-4}$ | 45.40% |
| | | $MAF \leq 1\times10^{-5}$ | 43.60% |
| | 0.8 | $0.005 < MAF < 0.01$ | 1.60% |
| | | $0.001 < MAF \leq 0.005$ | 5.90% |
| | | $1\times10^{-4} < MAF \leq 0.001$ | 14.30% |
| | | $1\times10^{-5} < MAF \leq 1\times10^{-4}$ | 44.40% |
| | | $MAF \leq 1\times10^{-5}$ | 33.80% |

Variants were obtained from chromosomes 1 and 2. The MAF for each variant is obtained from gnomAD for NFE. **Abbreviations**: ES, exome sequencing; gnomAD, Genome Aggregation Database; HRC, Haplotype Reference Consortium; MAF, minor allele frequency; NFE, non-Finnish European; $R^2$, imputation quality; and TOPMed, Trans-Omics for Precision Medicine.

**Supplementary Table 8D.** Variant Minor Allele Frequency Plot Statistics for Figure 1D

| Data | $R^2$ (>) | MAF category | Proportion |
|---|---|---|---|
| ES | - | $0.005 < MAF < 0.01$ | 0.20% |
| | | $0.001 < MAF \leq 0.005$ | 0.70% |
| | | $1\times10^{-4} < MAF \leq 0.001$ | 2.60% |
| | | $1\times10^{-5} < MAF \leq 1\times10^{-4}$ | 20.50% |
| | | $MAF \leq 1\times10^{-5}$ | 75.90% |
| HRC | 0.3 | $0.005 < MAF < 0.01$ | 3.40% |
| | | $0.001 < MAF \leq 0.005$ | 12.30% |
| | | $1\times10^{-4} < MAF \leq 0.001$ | 33.90% |
| | | $1\times10^{-5} < MAF \leq 1\times10^{-4}$ | 27.50% |
| | | $MAF \leq 1\times10^{-5}$ | 22.80% |
| | 0.8 | $0.005 < MAF < 0.01$ | 7.20% |
| | | $0.001 < MAF \leq 0.005$ | 20.10% |
| | | $1\times10^{-4} < MAF \leq 0.001$ | 32.30% |
| | | $1\times10^{-5} < MAF \leq 1\times10^{-4}$ | 20.60% |
| | | $MAF \leq 1\times10^{-5}$ | 19.80% |
| TOPMed | 0.3 | $0.005 < MAF < 0.01$ | 0.60% |
| | | $0.001 < MAF \leq 0.005$ | 2.10% |
| | | $1\times10^{-4} < MAF \leq 0.001$ | 7.80% |
| | | $1\times10^{-5} < MAF \leq 1\times10^{-4}$ | 39.60% |
| | | $MAF \leq 1\times10^{-5}$ | 49.90% |
| | 0.8 | $0.005 < MAF < 0.01$ | 1.60% |
| | | $0.001 < MAF \leq 0.005$ | 5.20% |
| | | $1\times10^{-4} < MAF \leq 0.001$ | 12% |
| | | $1\times10^{-5} < MAF \leq 1\times10^{-4}$ | 38.90% |
| | | $MAF \leq 1\times10^{-5}$ | 42.40% |
| HRC_TOPMed | 0.3 | $0.005 < MAF < 0.01$ | 0.60% |
| | | $0.001 < MAF \leq 0.005$ | 2.10% |
| | | $1\times10^{-4} < MAF \leq 0.001$ | 7.90% |
| | | $1\times10^{-5} < MAF \leq 1\times10^{-4}$ | 39.50% |
| | | $MAF \leq 1\times10^{-5}$ | 49.80% |
| | 0.8 | $0.005 < MAF < 0.01$ | 1.50% |
| | | $0.001 < MAF \leq 0.005$ | 5.20% |
| | | $1\times10^{-4} < MAF \leq 0.001$ | 13% |
| | | $1\times10^{-5} < MAF \leq 1\times10^{-4}$ | 38.90% |
| | | $MAF \leq 1\times10^{-5}$ | 41.40% |

Variants were obtained from chromosomes 1 and 2. The MAF for each variant is obtained from gnomAD for NFE. **Abbreviations**: ES, exome sequencing; gnomAD, Genome Aggregation Database; HRC, Haplotype Reference Consortium; MAF, minor allele frequency; NFE, non-Finnish European; $R^2$, imputation quality; and TOPMed, Trans-Omics for Precision Medicine.

**Supplementary Table 9A.** Variant Minor Allele Frequency Plot Statistics for Supplementary Figure 1A

| Data | $R^2$ (>) | MAF category | Proportion |
|---|---|---|---|
| ES | - | 0.005 < MAF < 0.01 | 0.10% |
| | | 0.001 < MAF ≤ 0.005 | 0.50% |
| | | $1\times10^{-4}$ < MAF ≤ 0.001 | 2.20% |
| | | $1\times10^{-5}$ < MAF ≤ $1\times10^{-4}$ | 21.30% |
| | | MAF ≤ $1\times10^{-5}$ | 75.80% |
| HRC | 0.3 | 0.005 < MAF < 0.01 | 3.60% |
| | | 0.001 < MAF ≤ 0.005 | 15% |
| | | $1\times10^{-4}$ < MAF ≤ 0.001 | 41% |
| | | $1\times10^{-5}$ < MAF ≤ $1\times10^{-4}$ | 29.50% |
| | | MAF ≤ $1\times10^{-5}$ | 10.90% |
| | 0.8 | 0.005 < MAF < 0.01 | 8% |
| | | 0.001 < MAF ≤ 0.005 | 29.50% |
| | | $1\times10^{-4}$ < MAF ≤ 0.001 | 39.90% |
| | | $1\times10^{-5}$ < MAF ≤ $1\times10^{-4}$ | 17% |
| | | MAF ≤ $1\times10^{-5}$ | 5.70% |
| TOPMed | 0.3 | 0.005 < MAF < 0.01 | 0.50% |
| | | 0.001 < MAF ≤ 0.005 | 1.90% |
| | | $1\times10^{-4}$ < MAF ≤ 0.001 | 7.50% |
| | | $1\times10^{-5}$ < MAF ≤ $1\times10^{-4}$ | 46% |
| | | MAF ≤ $1\times10^{-5}$ | 44.20% |
| | 0.8 | 0.005 < MAF < 0.01 | 1.40% |
| | | 0.001 < MAF ≤ 0.005 | 5.20% |
| | | $1\times10^{-4}$ < MAF ≤ 0.001 | 12.40% |
| | | $1\times10^{-5}$ < MAF ≤ $1\times10^{-4}$ | 46.10% |
| | | MAF ≤ $1\times10^{-5}$ | 34.90% |
| HRC_TOPMed | 0.3 | 0.005 < MAF < 0.01 | 0.50% |
| | | 0.001 < MAF ≤ 0.005 | 1.90% |
| | | $1\times10^{-4}$ < MAF ≤ 0.001 | 7.50% |
| | | $1\times10^{-5}$ < MAF ≤ $1\times10^{-4}$ | 46% |
| | | MAF ≤ $1\times10^{-5}$ | 44.10% |
| | 0.8 | 0.005 < MAF < 0.01 | 1.40% |
| | | 0.001 < MAF ≤ 0.005 | 5.30% |
| | | $1\times10^{-4}$ < MAF ≤ 0.001 | 13.40% |
| | | $1\times10^{-5}$ < MAF ≤ $1\times10^{-4}$ | 45.70% |
| | | MAF ≤ $1\times10^{-5}$ | 34.20% |

Variants were obtained from chromosomes 1 and 2. The MAF for each variant is obtained from gnomAD for NFE. **Abbreviations**: ES, exome sequencing; gnomAD, Genome Aggregation Database; HRC, Haplotype Reference Consortium; MAF, minor allele frequency; NFE, non-Finnish European; $R^2$, imputation quality; and TOPMed, Trans-Omics for Precision Medicine.

**Supplementary Table 9B.** Variant Minor Allele Frequency Plot Statistics for Supplementary Figure 1B

| Data | $R^2$ (>) | MAF category | Proportion |
|---|---|---|---|
| ES | - | 0.005 < MAF < 0.01 | 0.10% |
| | | 0.001 < MAF ≤ 0.005 | 0.40% |
| | | $1\times10^{-4}$ < MAF ≤ 0.001 | 1.80% |
| | | $1\times10^{-5}$ < MAF ≤ $1\times10^{-4}$ | 18.60% |
| | | MAF ≤ 1 $\times10^{-5}$ | 79.10% |
| HRC | 0.3 | 0.005 < MAF < 0.01 | 3.50% |
| | | 0.001 < MAF ≤ 0.005 | 13.50% |
| | | $1\times10^{-4}$ < MAF ≤ 0.001 | 40.30% |
| | | $1\times10^{-5}$ < MAF ≤ $1\times10^{-4}$ | 28.20% |
| | | MAF ≤ 1 $\times10^{-5}$ | 14.50% |
| | 0.8 | 0.005 < MAF < 0.01 | 7.60% |
| | | 0.001 < MAF ≤ 0.005 | 23.70% |
| | | $1\times10^{-4}$ < MAF ≤ 0.001 | 41.70% |
| | | $1\times10^{-5}$ < MAF ≤ $1\times10^{-4}$ | 19.50% |
| | | MAF ≤ 1 $\times10^{-5}$ | 7.60% |
| TOPMed | 0.3 | 0.005 < MAF < 0.01 | 0.40% |
| | | 0.001 < MAF ≤ 0.005 | 1.60% |
| | | $1\times10^{-4}$ < MAF ≤ 0.001 | 6.80% |
| | | $1\times10^{-5}$ < MAF ≤ $1\times10^{-4}$ | 42.70% |
| | | MAF ≤ 1 $\times10^{-5}$ | 48.50% |
| | 0.8 | 0.005 < MAF < 0.01 | 1.10% |
| | | 0.001 < MAF ≤ 0.005 | 4.30% |
| | | $1\times10^{-4}$ < MAF ≤ 0.001 | 11.80% |
| | | $1\times10^{-5}$ < MAF ≤ $1\times10^{-4}$ | 43.50% |
| | | MAF ≤ 1 $\times10^{-5}$ | 39.20% |
| HRC_TOPMed | 0.3 | 0.005 < MAF < 0.01 | 0.40% |
| | | 0.001 < MAF ≤ 0.005 | 1.60% |
| | | $1\times10^{-4}$ < MAF ≤ 0.001 | 6.90% |
| | | $1\times10^{-5}$ < MAF ≤ $1\times10^{-4}$ | 42.60% |
| | | MAF ≤ 1 $\times10^{-5}$ | 48.50% |
| | 0.8 | 0.005 < MAF < 0.01 | 1.10% |
| | | 0.001 < MAF ≤ 0.005 | 4.30% |
| | | $1\times10^{-4}$ < MAF ≤ 0.001 | 12.80% |
| | | $1\times10^{-5}$ < MAF ≤ $1\times10^{-4}$ | 43.30% |
| | | MAF ≤ 1 $\times10^{-5}$ | 38.50% |

Variants were obtained from chromosomes 1 and 2. The MAF for each variant is obtained from gnomAD for NFE. **Abbreviations**: ES, exome sequencing; gnomAD, Genome Aggregation Database; HRC, Haplotype Reference Consortium; MAF, minor allele frequency; NFE, non-Finnish European; $R^2$, imputation quality; and TOPMed, Trans-Omics for Precision Medicine.

**Supplementary Table 10.** Mean $R^2$ for all genes and significant genes for imputed datasets

| Dataset | Prevalence | MAF | μ $R^2$ (μ $R^2$) [a] |
|---|---|---|---|
| HRC | 0.1 | 0.01 | 0.723 (0.744) |
| | | 0.005 | 0.714 (0.734) |
| | | 0.001 | 0.682 (0.707) |
| | 0.2 | 0.01 | 0.723 (0.743) |
| | | 0.005 | 0.714 (0.734) |
| | | 0.001 | 0.682 (0.707) |
| TOPMed | 0.1 | 0.01 | 0.677 (0.682) |
| | | 0.005 | 0.675 (0.680) |
| | | 0.001 | 0.669 (0.673) |
| | 0.2 | 0.01 | 0.677 (0.681) |
| | | 0.005 | 0.675 (0.680) |
| | | 0.001 | 0.669 (0.672) |

Variants were obtained from chromosomes 1 and 2. All genes refer to genes with at least two variants. Significant genes refer to genes that met an exome-wide significance level of $2.5\times10^{-6}$ when simulation effect size = 1.5 and the proportion of causal rare variants with each gene region was 100%. Imputed variants with $R^2 > 0.3$ were retained. [a] Mean $R^2$ for all genes (Mean $R^2$ for significant genes). **Abbreviations**: μ, mean; HRC, Haplotype Reference Consortium; LoF, loss of function; MAF, minor allele frequency; $R^2$, imputation quality; and TOPMed, Trans-Omics for Precision Medicine.

**Supplementary Table 11.** Power to detect rare variant aggregate association and mean $R^2$ for overlapped gene sets for ES and TOPMed data

| Prevalence | MAF | Dataset | Power | μ $R^2$ |
|---|---|---|---|---|
| 0.1 | 0.01 | ES[a] | 0.716 | - |
| | | ES | 0.588 | - |
| | | TOPMed | 0.532 | 0.694 |
| | 0.005 | ES[a] | 0.675 | - |
| | | ES | 0.532 | - |
| | | TOPMed | 0.456 | 0.693 |
| | 0.001 | ES[a] | 0.531 | - |
| | | ES | 0.329 | - |
| | | TOPMed | 0.212 | 0.686 |
| 0.2 | 0.01 | ES[a] | 0.721 | - |
| | | ES | 0.582 | - |
| | | TOPMed | 0.531 | 0.694 |
| | 0.005 | ES[a] | 0.678 | - |
| | | ES | 0.524 | - |
| | | TOPMed | 0.460 | 0.693 |
| | 0.001 | ES[a] | 0.541 | - |
| | | ES | 0.326 | - |
| | | TOPMed | 0.217 | 0.686 |

Variants were obtained from chromosomes 1 and 2. Power was estimated as the number of genes that met an exome-wide significance level of $2.5\times10^{-6}$ divided by the maximum number of genes tested for ES, HRC, TOPMED, HRC_TOPMed, or ES_HRC_TOPMed, when simulation effect size = 1.5 and proportion of causal rare variants with each gene region was 100%. Imputed variants with $R^2 > 0.3$ were retained. [a] ES with all genes.
**Abbreviations**: μ, mean; ES, exome sequence; LoF, loss of function; MAF, minor allele frequency; $R^2$, imputation quality; and TOPMed, Trans-Omics for Precision Medicine.
